# Evaluation and comparison of methods for neuronal parameter optimization using the Neuroptimus software framework

**DOI:** 10.1101/2024.04.03.587881

**Authors:** Máté Mohácsi, Márk Patrik Török, Sára Sáray, Luca Tar, Szabolcs Káli

**Affiliations:** Institute of Experimental Medicine, Budapest, Hungary; Faculty of Information Technology and Bionics, Pázmány Péter Catholic University, Budapest, Hungary

**Author notes:** Corresponding author (SK).

## Abstract

Finding optimal parameters for detailed neuronal models is a ubiquitous challenge in neuroscientific research. Recently, manual model tuning has been replaced by automated parameter search using a variety of different tools and methods. However, using most of these software tools and choosing the most appropriate algorithm for a given optimization task require substantial technical expertise, which prevents the majority of researchers from using these methods effectively. To address these issues, we developed a generic platform (called Neuroptimus) that allows users to set up neural parameter optimization tasks via a graphical interface, and to solve these tasks using a wide selection of state-of-the-art parameter search methods implemented by five different Python packages. Neuroptimus also offers several features to support more advanced usage, including the ability to run most algorithms in parallel, which allows it to take advantage of high-performance computing architectures. We used the common interface provided by Neuroptimus to conduct a detailed comparison of more than twenty different algorithms (and implementations) on six distinct benchmarks that represent typical scenarios in neuronal parameter search. We quantified the performance of the algorithms in terms of the best solutions found and in terms of convergence speed. We identified several algorithms, including covariance matrix adaptation evolution strategy and particle swarm optimization, that consistently found good solutions in all of our use cases. By contrast, some other algorithms including all local search methods provided good solutions only for the simplest use cases, and failed completely on more complex problems. Finally, we created an online database that allows uploading, querying and analyzing the results of optimization runs performed by Neuroptimus, which enables all researchers to update and extend the current benchmarking study. The tools and analysis we provide should aid members of the neuroscience community to apply parameter search methods more effectively in their research.

**Author summary:** Model fitting is a widely used method in scientific research. It involves tuning the free parameters of a model until its output best matches the corresponding experimental data. Finding the optimal parameter combination can be a difficult task for more complex models with many unknown parameters, and a large variety of different approaches have been proposed to solve this problem. However, setting up a parameter search task and employing an efficient algorithm for its solution requires considerable technical expertise. We have developed a software framework that helps users solve this task, focusing on the domain of detailed models of single neurons. Our open-source software, called Neuroptimus, has a graphical interface that guides users through the steps of setting up a parameter optimization task, and allows them to select from more than twenty different algorithms to solve the problem. We have also compared the performance of these algorithms on a set of six parameter search tasks that are typical in neuroscience, and identified several algorithms that delivered consistently good performance. Finally, we designed and implemented a website that allows users to view and analyze our results and to add their own results to the database.

## Introduction

The construction and simulation of data-driven models has become a standard tool in neuroscience [1, 2, 3]. Such models can be employed, among other things, to consolidate the knowledge obtained from various experimental approaches into a common framework, to test the consistency of the data, and to make novel predictions by examining the response of the model to arbitrary inputs and by applying clean manipulations. Models at a given level of description (e.g., individual neurons) can also be combined to form models of entities at higher levels (such as networks) and thus aid the mechanistic understanding of emergent phenomena. Nevertheless, these data-driven models often contain parameters that are not directly constrained (or are only weakly constrained) by the available experimental data. Traditionally, such unknown parameters were often tuned manually to adjust the behavior of the model towards some desired target. However, this approach is typically inefficient, not quantitative, and may be heavily biased to reproduce a few selected experimental results at the expense of other relevant data. Consequently, in recent years, automated parameter search has emerged as the preferred method for the estimation of unknown parameters of neural models [4, 5, 6, 7, 8, 9, 10, 11, 12, 13, 14, 15, 16]. This approach requires the definition of an error function (or cost function) that measures the quality of the model with a given set of parameters, often in terms of how well it approximates data obtained using a particular experimental protocol. The goal of parameter optimization is then to find the set of parameters that minimizes the selected cost function. The difficulty of this task can vary widely depending on the nature and complexity of the model, the definition of the error function (or multiple error functions representing different goals, or objectives), and the number of unknown parameters. Simple optimization problems can be solved effectively by traditional gradient-based, local methods or by random search, but these approaches tend to fail when there are many unknown parameters and the cost function has multiple local minima [16, 17]. In fact, no algorithm is guaranteed to find the globally optimal parameter combination in a short time for all problems [18], and various clever search methods (called metaheuristics) have been proposed that often find good solutions in an acceptable amount of time by taking advantage of various types of regularities in the cost function [19].

Previous studies in neuroscience have used a variety of different software tools and algorithms to perform parameter optimization. The general-purpose neural simulators NEURON [20] and GENESIS [21] both include implementations of a few selected methods that are adequate for certain parameter search tasks. In addition, several tools have been developed specifically for neural parameter optimization, including Neurofitter [22], BluePyOpt [23], pypet [24], and NeuroTune [25], and some more general computational neuroscience tools such as NetPyNE [26] also have some support for parameter optimization. However, most of these tools rely on a very limited set of parameter search methods, which typically does not include many optimization algorithms that represent the state of the art in global optimization and are popular in other fields of science and engineering. These new methods were not included in any previous surveys of neural optimization. Systematic comparisons of the existing neural optimization software tools have also been quite limited [15]. Therefore, it is currently unknown which parameter search methods can be expected to perform well in the parameter optimization tasks that are typical in neuroscience.

Furthermore, most of the existing tools for neural optimization lack any intuitive user interface, and require substantial programming experience. One exception is our earlier optimization software called Optimizer [15], which included a graphical user interface (GUI) that was designed to guide users through the process of setting up, running, and evaluating the results of a neuronal parameter optimization task. Optimizer also provided four different optimization algorithms in two different Python packages, and was designed in a modular way to facilitate the integration of new components including additional optimization algorithms.

The goal of the current study was twofold. First, we aimed to provide a general software framework that allows the straightforward application of a large variety of state-of-the-art parameter optimization methods to typical problems in data-driven neural modeling. This was accomplished by significantly updating and extending our software tool (which is now called Neuroptimus). Second, we aimed to perform a systematic comparison of parameter search methods (including both previously used and novel algorithms) in the context of modeling single neurons, which is probably the most common subtype of parameter optimization tasks in neuroscience. To this end, we designed and implemented a test suite of neuronal parameter optimization problems, and used Neuroptimus to systematically test the performance of a large set of optimization algorithms on each of these benchmarks. The results of the different algorithms on the test suite were systematically analyzed and compared. Finally, we designed and deployed a web-accessible database that contains all the results of this study and also allows users to upload, retrieve, and analyze the results of parameter optimization.

## Results

The systematic evaluation of parameter optimization methods in the context of neuronal modeling required the development of several interrelated methods and tools, which are described in detail in the Methods section and whose main features are also summarized below. The first necessary ingredient was a software tool that allows users to set up, execute, and evaluate the results of a wide variety of neural parameter optimization problems in a single standardized framework. The second required component was a diverse set of benchmark problems that differ in the type of the model, the number of unknown parameters, and the complexity of the error function, and that collectively cover many types of parameter fitting problems that are often encountered in neuronal modeling. The third necessary component was a set of methods that allows the consistent evaluation and comparison of optimization results across the different benchmarks and algorithms. Finally, the last ingredient was a web-accessible database of the optimization results that allows us to share all of our results publicly and also enables us as well as other researchers to extend the study with additional optimization runs and even new benchmarks.

### The Neural Optimization User Interface (Neuroptimus)

We began our study by updating, improving and extending our previously developed optimization software (Optimizer), which was already shown to be a useful tool for neuronal optimization [15]. The new version (named Neuroptimus) inherited many useful features from its predecessor, and added several important new capabilities. Both Optimizer and Neuroptimus support the definition and solution of neural optimization problems through a graphical user interface (GUI) that guides the users throughout the process. The main steps (represented by different tabs in the GUI) involve selecting the target data, selecting the model and the parameters to be optimized, setting up the simulations (including stimulation and recording parameters), defining the cost function, selecting the optimization algorithm, running the parameter search, and reviewing the results. A detailed guide to the GUI is available in the online documentation of Neuroptimus (https://neuroptimus.readthedocs.io/en/latest/). All the functionality is also accessible through a command line interface that uses configuration files to set up the optimization, which enables batch processing (e.g., multiple runs with different settings or random seeds). Simulations of the model can be performed either by the NEURON simulator [20] (which is handled internally) or by arbitrary external code (which may include running other simulators) handled as a “black box”. The modular, object-oriented structure of the program makes it possible to extend its capabilities by adding new error functions and optimization algorithms.

Neuroptimus includes several new and enhanced features compared to Optimizer. In addition to specific time series (such as voltage traces), it is now also possible to use as target data the statistics of features extracted (e.g., using the feature extraction module eFEL, [27]) from a set of experimental recordings. In this case, Neuroptimus uses eFEL to extract the same features from each simulated model, computes feature errors as the difference between the feature value of the model and the mean value of the experimental feature, normalized by the experimental standard deviation, and uses the sum of these feature errors as the cost function during parameter optimization. Weights can also be provided individually for each error component.

While Optimizer provided four different search algorithms (two local and two global algorithms implemented by the Inspyred and Scipy packages), Neuroptimus currently supports more than twenty different optimization algorithms from five external Python packages (see Table 2 for a complete list), plus an internally implemented random sampling algorithm, which can be considered as a simple baseline method.

Neuroptimus also contains many enhancements “under the hood”. The new version was entirely developed in Python 3 to support recent open-source Python modules, such as search algorithms, graphical and parallelization interfaces. The graphical user interface was completely re-implemented using the PyQt5 package, which provides a Python binding to the popular cross-platform GUI toolkit Qt. In addition to the parameter search methods offered by Scipy and Inspyred, Neuroptimus now also provides an interface to the algorithms implemented by the widely used optimization packages Pygmo and BluePyOpt, as well as an additional parallelized Python implementation of the Covariance Matrix Adaptation Evolution Strategy (CMA-ES) algorithm. For many of these search algorithms, parallel evaluation of models is also supported and easily configurable, which can lead to a manifold reduction in execution time, especially on highly parallel architectures such as compute clusters and supercomputers.

### Neural optimization benchmarks

We defined and implemented a test suite of different neuronal optimization problems to demonstrate the utility of our Neuroptimus software and to quantitatively evaluate and compare the effectiveness of different parameter optimization algorithms. Our aim was to identify which parameter search methods (and which implementations) are able to find good solutions to each of our benchmark problems, and which methods (if any) can provide consistently good performance across all of these tasks. Our benchmarking use-cases differ in the complexity of the models, the simulation protocol, the source and nature of the target data, the features and error functions used to evaluate the model, and the number of unknown parameters. A subset of our use-cases is analogous to those that were described by Friedrich et al. [15], although some of these have been updated to improve their robustness. Each of the six benchmark problems is described briefly below, and in more detail in the Methods section.

Four of the use cases involve finding the biophysical parameters of compartmental models of neurons based on somatic voltage responses; however, these models differ greatly in terms of the level of morphological and biophysical detail, and also in the number of unknown parameters (between 3 and 12). One simple use case involves the classic single-compartment Hodgkin-Huxley model with two voltage-gated conductances and a leak conductance; one uses a morphologically detailed but passive model neuron; another benchmark optimizes the somatic conductances of several voltage-gated ion channels in a simplified (6-compartment) model, while our most complex use case involves fitting spatially varying conductance densities for a large set of ion channels in a fully detailed compartmental model of a hippocampal pyramidal cell. A different type of benchmark involves optimizing the parameters of a phenomenological point neuron (an adaptive exponential integrate-and-fire model), and the final one simulates a voltage-clamp experiment to estimate synaptic parameters.

Some of our benchmark problems (the Hodgkin-Huxley and the Voltage Clamp use-cases) use surrogate data as the target. In this case, target data are generated by the same neuronal model with known parameters; some of these parameters are then considered to be unknown, and the task is to reconstruct the correct values. Therefore, in these test cases, a perfect solution with zero error is known to exist, and the corresponding parameters can be compared to those found by the search algorithms. However, for most of our benchmark problems, the target data were recorded in electrophysiological experiments, or (in one case) generated by a more complex model than the one we were fitting. In these instances, the best-fitting parameters and the minimal possible error score are unknown.

In most of our use cases we compared the output of the model to the target data by extracting several different electrophysiological features from the raw voltage traces. The difference of each model feature from the corresponding (mean) experimental feature can be considered as a separate error component (or objective). This allowed the direct application of multi-objective optimization methods. When using single-objective algorithms, feature errors were combined into a single cost function using an average with pre-defined (in most cases, uniform) weights. The final best solution for multi-objective algorithms was also chosen using the same weighted average of the objectives. Two of our use cases had simpler voltage or current traces as their target. In these cases, the mean squared difference between the model trace and the experimental trace was used as the only error function. This precluded the use of multi-objective optimization methods, so only single-objective algorithms were included in the comparison in these cases.

We used several criteria to select optimization algorithms for inclusion in our benchmark study. First, we implemented a simple random search algorithm based on independently repeated uniform sampling of the entire available search space defined by the parameter boundaries. This algorithm can be considered as a natural baseline against which we can measure the performance of more sophisticated methods. Second, we included some popular local optimization algorithms (Nelder-Mead and L-BFGS-B) that are expected to be efficient when the error function has a single optimum, but not for more complex problems with multiple local optima. The rest of the search algorithms that we included are so-called global optimization methods or meta-heuristics, which aim to take advantage of certain types of regularities in the error function to find the global optimum (or another similarly good solution) more efficiently than a random search or local optimization methods do. A very large selection of such meta-heuristic algorithms has been developed, and many of these are included in one (or several) of the Python packages that are accessible in Neuroptimus. Due to time and resource constraints, not all of these algorithms were included in the current study, but we aimed to include many of the algorithms that were previously used in neuronal optimization and those that have proved particularly successful in other settings. More specifically, we included several different types of evolutionary algorithms, several implementations of the particle swarm algorithm, and also some other types of bioinspired algorithms and methods based on statistical physics.

To ensure a fair comparison of different search methods, we allowed a maximum of 10,000 model evaluations in a single run of every optimization algorithm. For all the algorithms that define populations of models that are evaluated as a batch in every iteration (this includes both evolutionary and swarm algorithms), we set the population size to 100, and ran the algorithms for 100 iterations (generations). We recorded the lowest error value achieved during each run, and also looked at how the best error score evolved during the course of the optimization. This allowed us to quantify the speed of convergence by calculating the area under the curve showing the cumulative minimum error as a function of completed model evaluations. We performed 10 repeated runs of each algorithm on every benchmark problem to allow proper statistical evaluation of the results.

### The performance of different optimization algorithms on individual benchmarks

#### Hodgkin-Huxley model

Our first benchmark problem involved finding the correct densities of two voltage-gated conductances and a leak conductance (3 parameters overall) in the classic single-compartment Hodgkin-Huxley model [28] based on the voltage response to a single current step stimulus (Figure 1). We compared the response of each candidate model to that of the original model by evaluating four features (spike count, spike amplitude, spike width, and mean squared error of the voltage excluding spikes, evaluated using built-in error functions of Neuroptimus), which also enabled the application of multi-objective optimization methods. We expected this to be a relatively simple optimization problem based on the low number of parameters to fit, although it is also clearly non-trivial due to the nonlinear nature of the neuronal dynamics and, particularly, the complicated dependence of the extracted physiological features on the conductance parameters.

**Figure 1.**
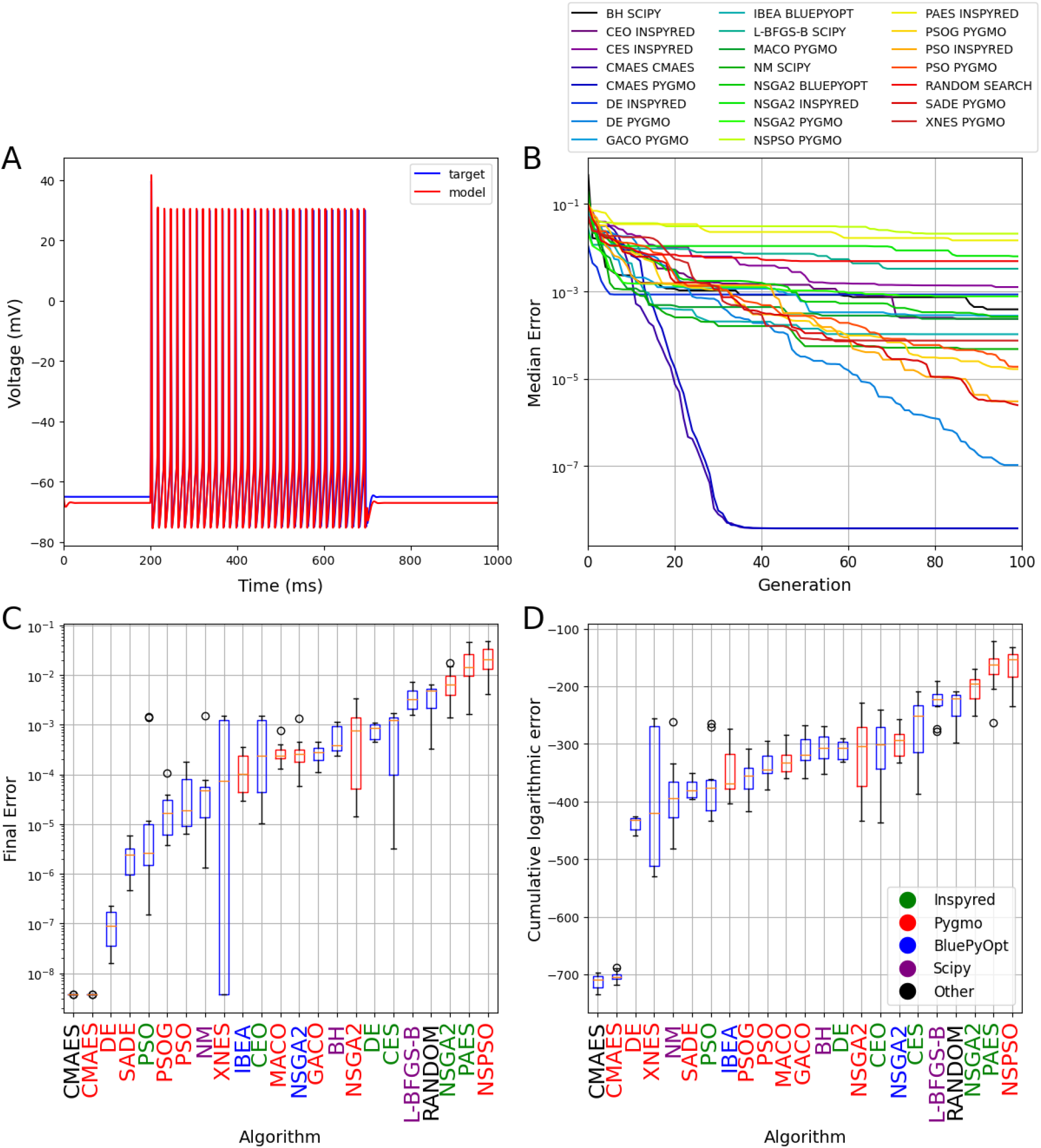
The results of fitting conductance densities in the Hodgkin-Huxley model. (A) Example of a comparison plot showing the voltage trace generated by the model with its original parameters (blue) and the trace given by the model using the best parameter set found by the Random Search algorithm (red). (B) Plot showing the evolution of the cumulative minimum error during the optimization. The curves show the median of 10 independent runs for each relevant algorithm. Each generation corresponds to 100 model evaluations. The colors corresponding to the different algorithms (and packages) are shown in the legend. (C) Box plot representing the distribution of the final error scores over 10 independent runs of each algorithm. (D) Box plot representing the convergence speed of the algorithms tested, measured as the area under the logarithmic cumulative minimum error curve (as shown in panel B). In (C) and (D), horizontal red lines indicate the median, the boxes represent the interquartile range, whiskers show the full range (excluding outliers), and circles represent outliers. Boxes representing single-objective algorithms are colored blue and those of multi-objective ones are red. Results are sorted by the median score, from the best to the worst. The names of the packages on the horizontal axis are colored to indicate the implementing package according to the legend in (D).

Many of the search algorithms tested found relatively good solutions most of the time, but most of them failed to converge completely in 10,000 model evaluations. The exception was the CMAES algorithm, whose implementations both consistently converged to the optimal solution after approximately 3,500 evaluations (the lowest possible error score was not exactly zero due to rounding errors). Interestingly, multi-objective algorithms generally performed worse on this use-case than single-objective ones, with Inspyred’s NSGA2, PAES and Pygmo’s NSPSO algorithms giving worse results than Random Search. Different implementations of the same algorithms (two versions for CMAES, three for PSO, and three for NSGA2) usually showed similar convergence behavior, except for the implementation of NSGA2 by the Inspyred package that performed significantly worse than the Pygmo and BluePyOpt versions of the same method. Overall, even this simple benchmark revealed surprisingly large differences in the performance of the different search methods that we included in our comparison.

#### Voltage Clamp

The second benchmark problem involved finding four parameters of a simulated synaptic connection to a single-compartment model neuron using voltage-clamp recordings (Figure 2). This use case also used surrogate data as the target, but in this case the recorded variable was the current injected by the electrode during a simulated voltage-clamp experiment. The parameters to be reconstructed were the maximal value (weight), delay, and rise and decay times of the synaptic conductance change following each repeated activation of the synapse. Due to the stereotyped nature of the data, mean squared difference was used as the only error function, and thus only single-objective algorithms were tested.

**Figure 2.**
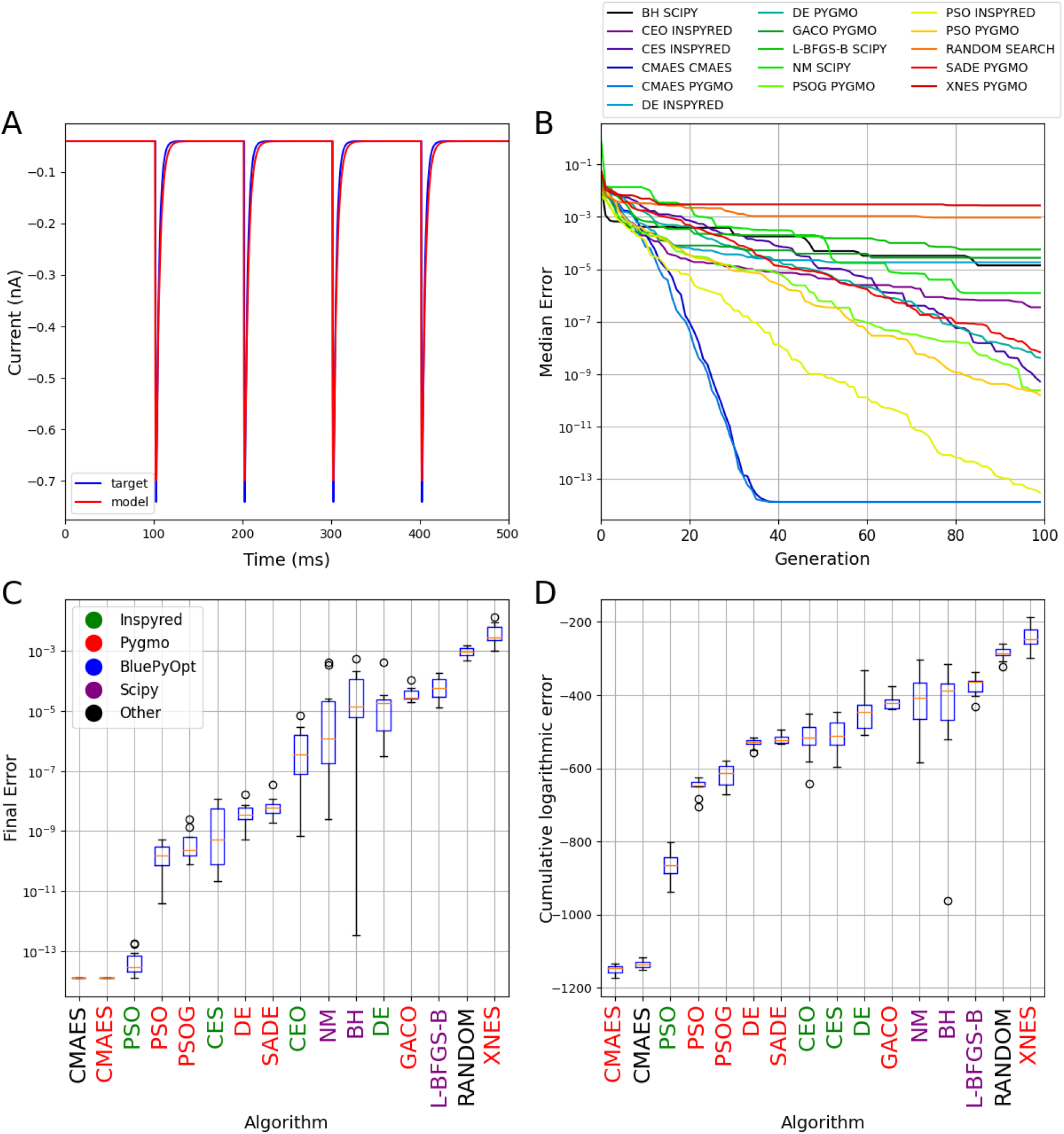
The results of fitting the parameters of a synaptic connection based on simulated voltage-clamp recordings. The plots in all four panels are analogous to those in Figure 1. Only single-objective methods were tested in this use-case because only a single error function (mean squared difference) was used to compare model outputs to the target data. Panel A shows the results of a best-fitting model found by the Random Search algorithm.

Although this is still a relatively simple and low-dimensional problem, and the intrinsic dynamics is much less complex than that of the Hodgkin-Huxley model in current clamp mode in the first use-case above, we observed highly divergent performance for the set of algorithms that we tested. Both implementations of CMAES reached the best possible score (again defined by round-off error) in fewer than 40 generations (4000 model evaluations). The Inspyred implementation of PSO also approached this limit by the end of the optimization (10,000 model evaluations), but it converged substantially slower than CMAES. The Pygmo implementations of PSO, two versions of the DE algorithm, and the CES algorithm of Inspyred also achieved good results, but converged even more slowly. At the other end of the spectrum, local search algorithms were typically not effective at solving this problem, and the XNES algorithm from the Pygmo package actually performed worse than the baseline random search method.

#### Passive, anatomically detailed neuron

This use-case represents an important practical problem that has been investigated in several previous studies [15, 29, 30, 31]. It involves the estimation of three basic biophysical parameters that determine the propagation and integration of voltage signals within neurons in the subthreshold voltage range: the (specific) membrane capacitance, membrane resistance, and axial resistance. The task is to estimate these three parameters based on the voltage response of a neuron to a current stimulus (which, in our case, consisted of a larger short and a smaller long current step) recorded from a hippocampal pyramidal cell in vitro (Figure 3). The response of the model is linear in terms of the injected current, but still depends on the combination of the three biophysical parameters (which are assumed to be spatially uniform within the cell) in a non-trivial way due to the complex morphology of the neuron. In the absence of spikes, we used the mean squared difference between the simulated and the experimentally recorded voltage traces as the only error function, and restricted our attention to single-objective algorithms.

**Figure 3.**
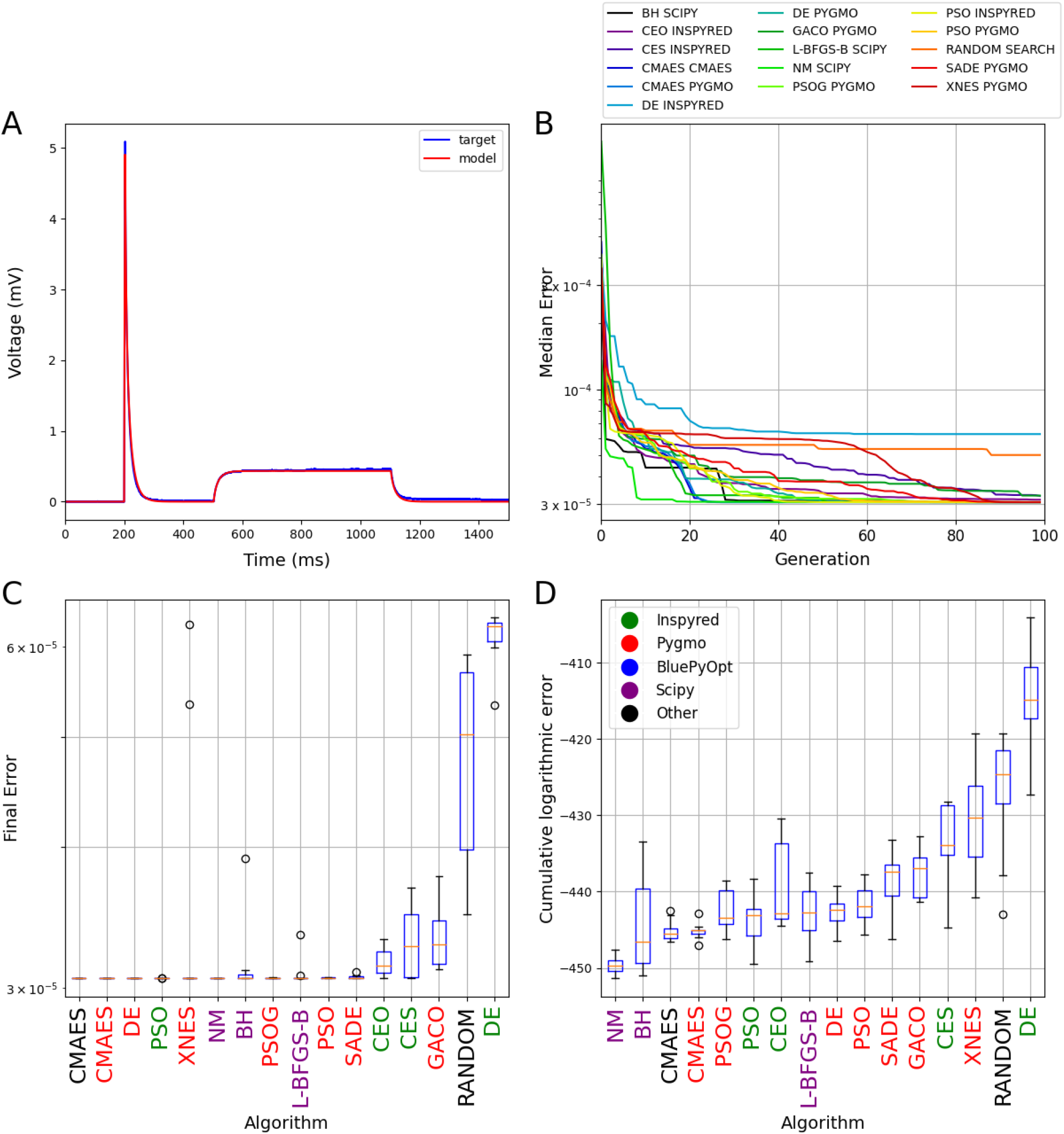
The results of fitting the passive biophysical parameters of a morphologically detailed multi-compartmental model to experimental recordings from a hippocampal pyramidal neuron. The plots in all four panels are analogous to those in Figure 1. Only single-objective methods were tested in this use-case because only a single error function (mean squared difference) was used to compare model outputs to the target data. Panel A shows the results of a best-fitting model found by the CMAES algorithm.

This benchmark proved to be the easiest in our entire test suite. Many algorithms found the best possible fit to the data in (almost) all the runs, and most of them also converged relatively rapidly. In this case, local search methods such as the Nelder-Mead and the L-BFGS-B algorithms also found the optimal solution efficiently in most runs. One curious exception was the DE algorithm implemented by the Inspyred package, which achieved a worse result than Random Search, even though the other implementation of the same algorithm by the Pygmo package was among the high-performing methods.

#### Simplified active model

This benchmark problem is more complex than the previous ones in several respects. The task in this use case is to determine the somatic densities of nine voltage-gated conductances in a model of a hippocampal CA1 pyramidal neuron with simplified morphology (consisting of only six compartments) so that the somatic voltage response of the model best approximates the response of a fully detailed CA1 pyramidal cell model under the same conditions (Figure 4). We used five of the error functions implemented by Neuroptimus (mean squared error excluding spikes, spike count, latency to first spike, action potential amplitude, action potential width, and after-hyperpolarization depth) to compare the two voltage traces. This also enabled us to test multi-objective algorithms besides the single-objective ones.

**Figure 4.**
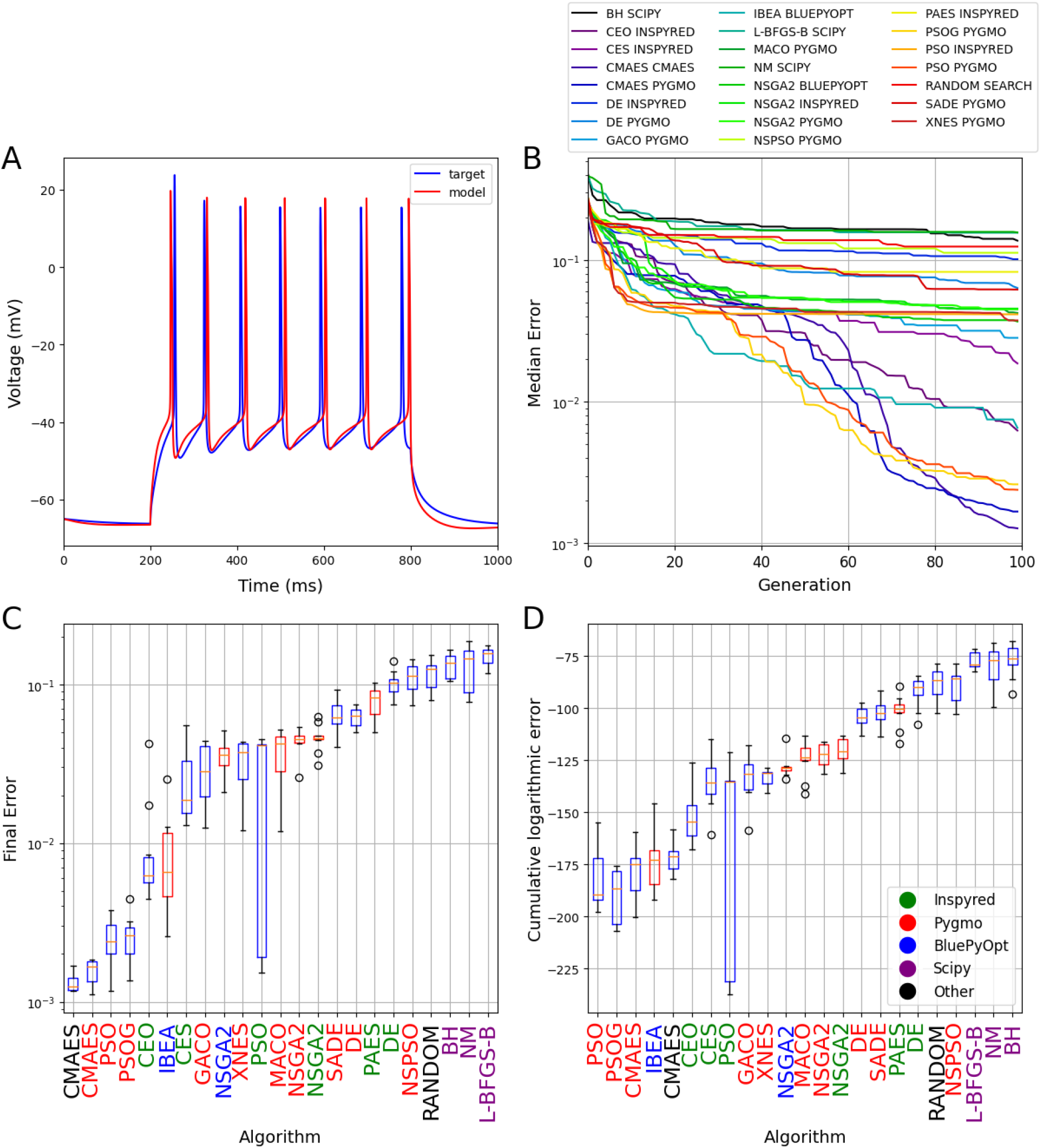
The results of fitting the densities of somatic voltage-gated conductances in a morphologically simplified six-compartment model using a simulated voltage trace from a detailed compartmental model as the target. The plots in all four panels are analogous to those in Figure 1. Panel A shows the results of a best-fitting model found by the CMAES algorithm.

In this more complex use case, there were large differences in performance among the algorithms, with two orders of magnitude difference between the final errors of the best- and the worst-performing methods. Once again, implementations of the CMAES algorithm achieved the best final scores, but the Pygmo implementations of PSO also delivered good final scores along with the best convergence speed. Among multi-objective algorithms, IBEA achieved the best final scores, and also performed quite well in terms of convergence speed. At the other extreme, all local search algorithms typically performed worse than Random Search, and are clearly inadequate for this type of problem. It is worth noting that all three implementations of the NSGA2 algorithm gave similar results, as did the different flavors of DE, although neither these algorithms nor several other bio-inspired algorithms (such as other evolutionary algorithms or ant colony optimization) were capable of providing as good solutions as CMAES and PSO on this benchmark.

#### Extended integrate-and-fire model

This use case involves fitting the parameters of an adaptive exponential integrate-and-fire model neuron so that it captures the spiking responses of a hippocampal CA3 pyramidal neuron recorded in vitro (Figure 5). This is a single-compartment model that does not include detailed models of neuronal biophysics; instead, it aims to capture neuronal spiking phenomenologically, using an extended integrate-and-fire formalism with an exponential term in the current-voltage relationship and an adaptation variable that is also linked to spiking [32, 33]. This model has a total of 10 parameters that had to be fitted by the optimization algorithms. Unlike the models in the other use cases (which were implemented in NEURON), this model was implemented in the NEST simulator [34], and was treated as a black box by Neuroptimus. The parameters generated by the optimization algorithms were passed to an external Python script, which constructed the model, ran the simulations using NEST, and passed the results (spike times and subthreshold voltage traces in two separate files) back to Neuroptimus for evaluation and comparison with the experimental data. The data included the voltage responses of a real CA3 pyramidal cell to current steps of four different amplitudes (these responses are shown concatenated in blue in Figure 5A), and the model had to capture all of these responses simultaneously. As integrate-and-fire models cannot (and are not expected to) reproduce spike shape, we used spike count, latency to first spike, and the mean squared difference of the voltage excluding spikes as three error components during the optimization.

**Figure 5.**
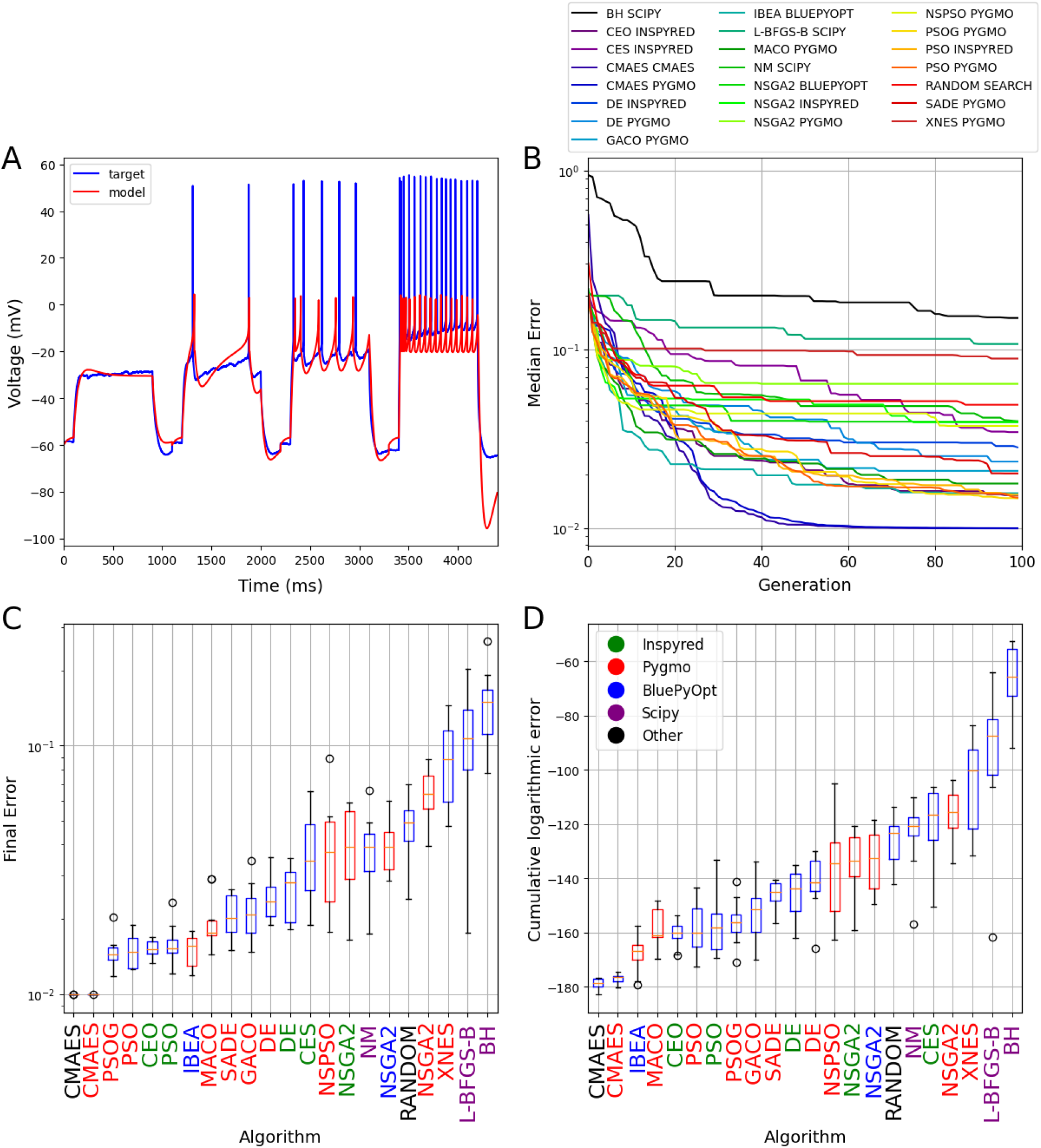
The results of fitting a phenomenological spiking neuronal model (the adaptive exponential integrate-and-fire model) to capture experimental recordings with multiple traces. The plots in all four panels are analogous to those in Figure 1. Panel A shows the results of a best-fitting model found by the CMAES algorithm. Note that the height of action potentials is irrelevant in the integrate-and-fire model, and the spikes generated by the model are not explicitly represented in the figure.

On this benchmark, the two implementations of CMAES found the solutions with the lowest error. In fact, they obtained the same lowest error score several times, and this was lower than the scores achieved by any other algorithm, so this error score likely corresponds to the best possible solution of this optimization problem. Although clearly inferior to CMAES on this problem, the various implementations of the particle swarm algorithm, the multi-objective algorithm IBEA, and the classical evolutionary algorithm found relatively good solutions, while several methods performed substantially worse than Random Search. We note that the PAES algorithm generated parameter combinations that led to errors during the NEST simulation, and was therefore excluded from the current comparison.

#### Morphologically and biophysically detailed CA1 pyramidal cell model

Our final use case represents a typical scenario in the construction of morphologically and biophysically detailed compartmental models [1, 4, 7, 35, 36, 37, 38]. The model is based on the reconstructed morphology of a CA1 pyramidal neuron [39], and contains a large set of voltage-gated conductances, several of which are distributed non-uniformly within the cell (see Methods for further details of the model). The goal is to find the values of 12 parameters that determine the densities and biophysical properties of voltage-gated and leak conductances in the model such that the features extracted from the voltage responses of the model to multiple step current injections best approximate the average of the same features extracted from experimental recordings under matching conditions (Figure 6). One hyperpolarizing and five depolarizing current steps were used, and these yielded a total of 66 features of 20 different types (Table 3) that were extracted and evaluated for each model instance during the parameter search.

**Figure 6.**
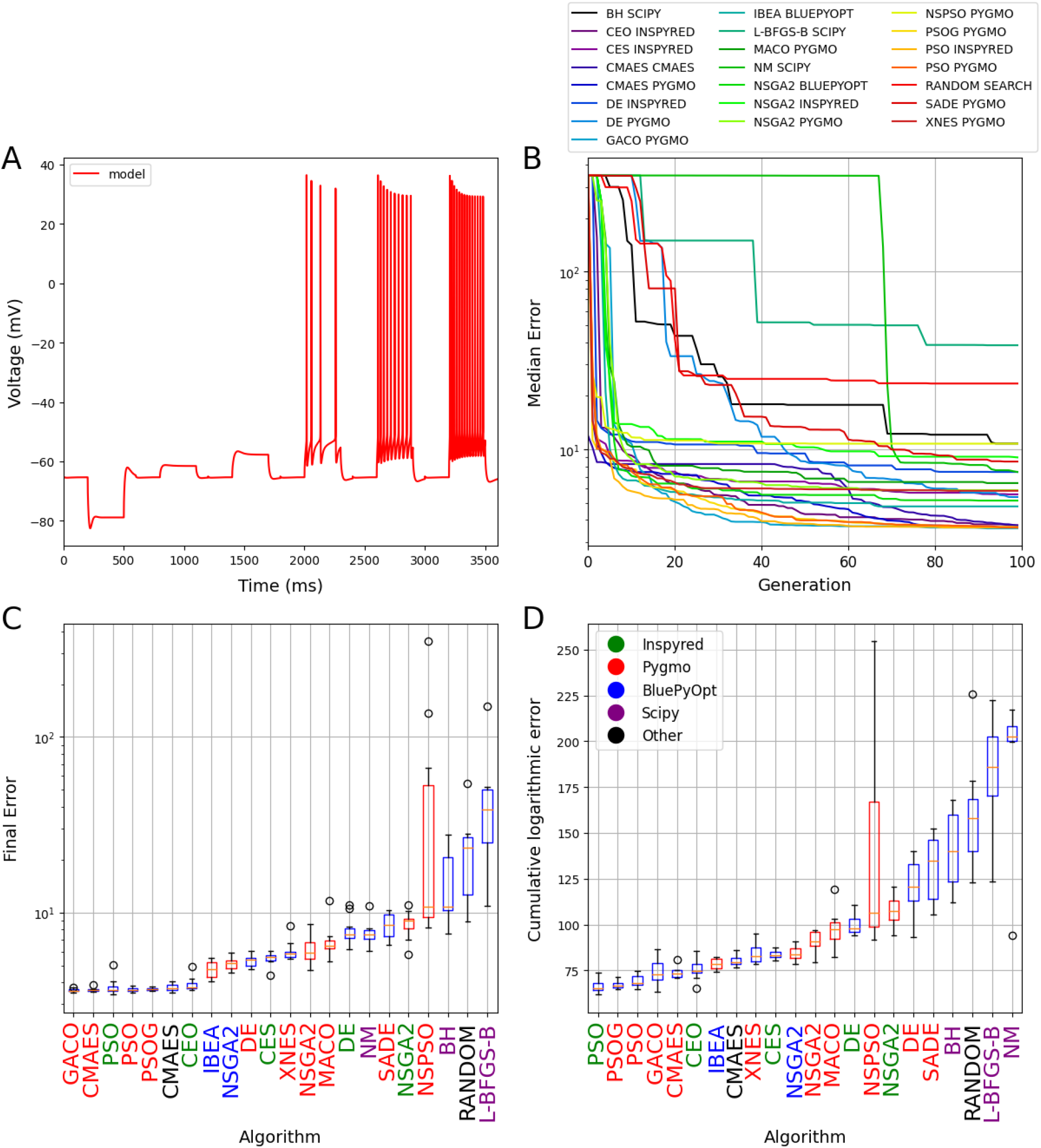
The results of fitting conductance densities and kinetic parameters in a CA1 pyramidal cell model. The plots in all four panels are analogous to those in Figure 1. Panel A shows the results of a best-fitting model found by the CMAES algorithm. No target trace is shown because, in this use case, the actual target is defined by the statistics of electrophysiological features that are extracted from a set of experimental recordings.

Although this is certainly the most complex model in our benchmarking suite with the largest number of free parameters, finding solutions with errors close to the smallest possible value was apparently easier than in the previous two use-cases (although, strictly speaking, we cannot rule out the possibility that none of the algorithms tested ever came close to the unknown globally optimum error score). More specifically, all three versions of PSO, both implementations of CMAES, and also the GACO and CEO algorithms consistently yielded similar low error scores, but several other algorithms, including the multi-objective IBEA and NSGA2 methods, also gave acceptable solutions. We note that running the PAES algorithm resulted in memory errors, and it was therefore omitted from the evaluation of this use case.

### Overall performance of the algorithms

In general, no single algorithm is expected to perform well in all types of global optimization problems. Popular methods can take advantage of different types of regularities in the error function to speed up the search for the global optimum even in high-dimensional spaces with multiple local optima. Therefore, problems with different structures may require different algorithms for their efficient solution, and we can identify some signs of this heterogeneity when comparing the results of the individual benchmarks described above. Nevertheless, some clear patterns are evident, and we can quantify this by constructing and examining summary statistics for the algorithms across all the use cases.

XXFigure 7 summarizes the rankings of the various algorithms in our study according to final score and convergence speed. Individual ranks were based on the medians of the respective performance measure across all runs of the algorithm in a particular benchmark (this was also the basis of the placement of the algorithms along the horizontal axes in panels C and D of Figures 1-6), and Figure 7 shows the statistics of these ranks for each algorithm across the different benchmarks.

**Figure 7.**
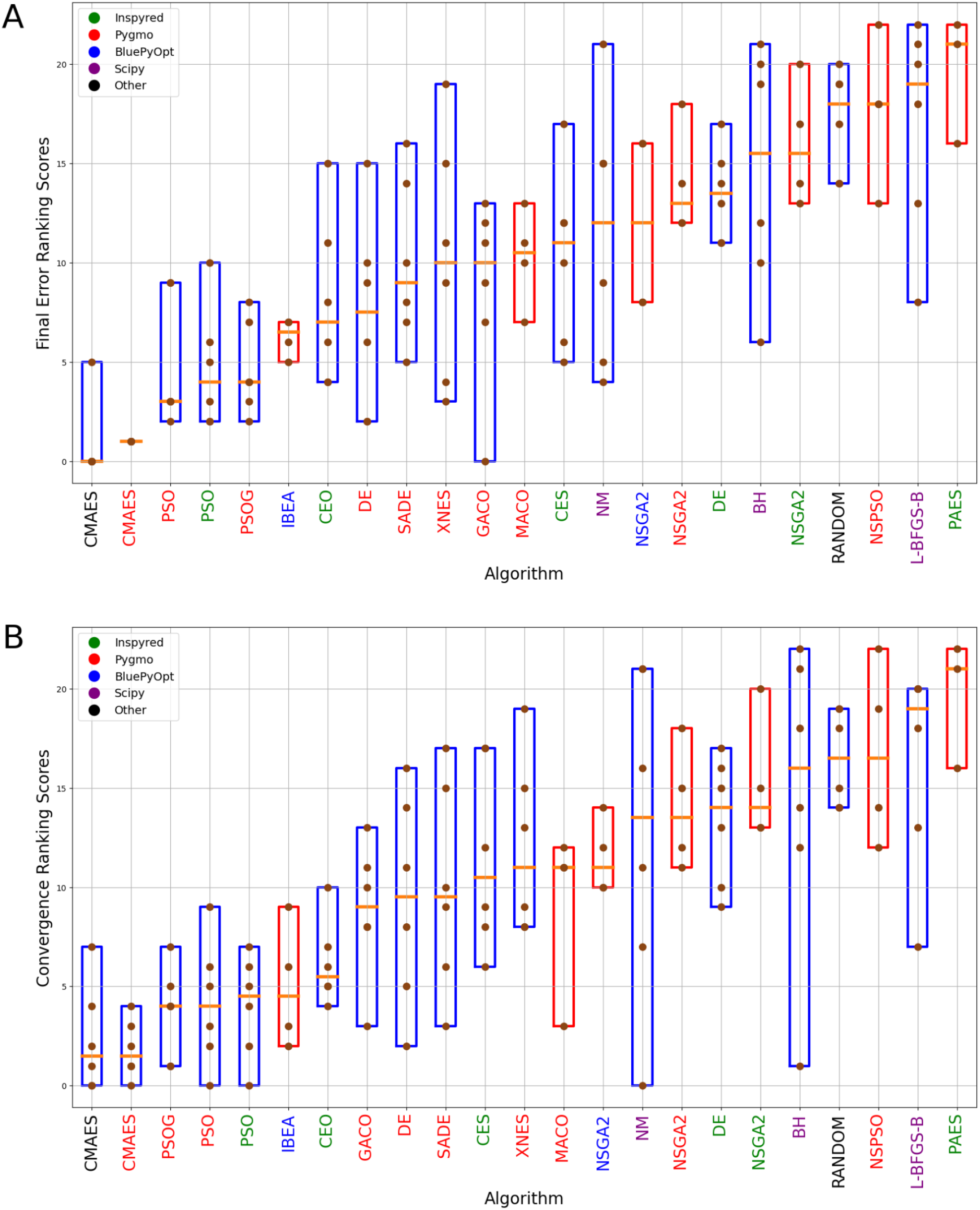
Overall rankings of optimization algorithms. Statistics of the ranks achieved by individual optimization algorithms on the different benchmarks (Figures 1-6) according to the final error (A) and convergence speed (B). Brown dots represent the ranks achieved by the algorithms in each use-case; boxes indicate the full range and the orange line represents the median of these ranks. The single-objective algorithms are shown in blue and the multi-objective ones in red boxes. The color of the name of the algorithm indicates the implementing package, with the color code included in the legend. Algorithms are sorted according to the median of their ranks.

In terms of the generally best-performing algorithms on our neuronal optimization test suite, the results are quite clear. In almost all cases, CMAES delivered the best results after 10,000 model evaluations, and its two implementations by different packages performed quite similarly. The three implementations of the particle swarm algorithm that we tested also showed similar performance, and were typically better than all the other methods except for CMAES. IBEA was close behind the PSO variants in the rankings, and was clearly the best among the multi-objective methods that we tested. It is interesting to note that some of the algorithms, including local search methods (and especially the Nelder-Mead algorithm) but also some other methods such as GACO and XNES showed widely varying performance across the different benchmarks, so these may be suitable for some problems but completely inadequate for others. Finally, the rankings based on the convergence score are generally quite similar to those based on just the final score, although there are some minor differences - for instance, PSO appears to be more competitive with CMAES according to this measure.

### Online database of optimization results

The results presented so far summarize the performance of a selected subset of the algorithms implemented by five Python packages, using their default settings, on a pre-defined suite of six neuronal optimization problems. To increase the utility and reproducibility of our results, we also wanted to share the details of all the optimization runs, including the settings that enable their replication as well as their detailed results. In addition, we wanted to find a simple way of updating and extending the study with more optimization runs, potentially with different settings or algorithms not included in the present comparison, or even involving additional use cases, not just by us but also other interested researchers. We therefore designed, implemented and deployed an online database with an associated, publicly accessible web server (https://neuroptimus.koki.hu) that allows users to upload, query, and analyze optimization runs performed by the Neuroptimus software tool.

The website allows users to browse the optimization results stored in the online database, and filtering options are available to create lists of relevant results. The results of optimization runs can be viewed in a detailed text-based format, and selected subsets of optimizations can be analyzed and displayed graphically, similarly to the plots in Figures 1-6 above. Registered users can also add to the database their own optimization results by uploading the JSON file (metadata.json) generated by Neuroptimus after each optimization run. Users can optionally also upload the other files that belong to the optimization (including the model and the target data) in the form of an archive, which creates an online record of the optimization that allows its full replication.

The database currently contains the results of all the optimization runs from the current study. This enables users of the website to replicate most of the figures in this paper, and to download individual optimization runs (including their settings and results). Users can also carry out custom analyses of the results, and (after registration) they can add their own optimization results (created using Neuroptimus) on both existing and novel use cases and compare these with other results on the same use case. This way, the website offers an interactive, continuously updated, and publicly accessible “live” version of this paper, which will provide a valuable online resource for researchers to explore and share methods and results on neuronal optimization.

## Discussion

The results of our study have the potential to advance the state of the art in neural parameter optimization in several different ways. First, we have created and shared Neuroptimus, a software tool that was designed to help both computational and experimental neuroscientists in the complete formulation and solution of neuronal parameter search problems. All the functions of Neuroptimus are accessible through a graphical user interface, although there is also a command line interface to support more advanced usage. Users of Neuroptimus gain uniform access to a large number of optimization algorithms implemented by several widely used Python packages, including several algorithms that were used successfully in previous neuronal modeling studies, and also several other state-of-the-art optimization methods that are popular in other domains but have not been applied to neuronal parameter optimization. This feature of Neuroptimus allowed us to systematically test the performance of a wide variety of parameter search methods on six distinct neuronal optimization problems, which makes it possible to offer some recommendations for future neuroscientific studies that rely on parameter optimization (see below). Finally, we created an online database of optimization results obtained by using Neuroptimus. This database currently contains the results of the present study, but the online user interface also allows us as well as others to add new results and compare them with already existing ones.

### Comparison of Neuroptimus with other neural optimization tools

A variety of software tools have been developed and used for the purpose of optimizing the parameters of neural models. This includes the built-in optimization modules of the general-purpose neural simulators NEURON [20] and GENESIS [21], the optimization-oriented features of the NetPyNE neural modeling framework [26], the Neurofitter program [22], as well as the Python packages BluePyOpt [23], NeuroTune [25], and pypet [24]. However, each of these tools (except for Neurofitter, which is no longer actively maintained, and focuses on a single specific cost function) relies on just one or a few algorithms, or a single external optimization package, to perform parameter search, while Neuroptimus provides access to a large variety of different algorithms from five distinct Python packages. This gives users of Neuroptimus a lot of flexibility to choose the best method for any particular fitting problem. As an example, Neuroptimus can take advantage of the large number of local and global optimization algorithms offered by the Pygmo package [40], which is a mature and actively maintained tool used, among others, by the European Space Agency. Another distinguishing feature of Neuroptimus is its graphical user interface. Among the other tools, only NEURON offers GUI-based access to parameter optimization, but the utility of this feature is severely limited by its reliance on the local search method PRAXIS as its only available algorithm.

### Algorithm recommendations based on our benchmarking results

The performance of optimization algorithms in general depends quite heavily on the nature of the problem, and no particular algorithm is expected to provide good solutions universally. However, within the task domain that we considered here, i.e., finding the biophysical parameters of models of single neurons, we can make some clear recommendations. Our benchmarking results were dominated by two different metaheuristics, covariance matrix adaptation evolution strategy (CMAES) and particle swarm optimization (PSO), followed by the multi-objective indicator-based evolutionary algorithm (IBEA) - so we would suggest trying these methods first to attack a novel neuronal parameter optimization problem. We also confirmed that local optimization algorithms are generally not suitable for more complex parameter search tasks, although they can be adequate and even efficient in the simplest cases.

Although, in principle, implementation details (particularly the default settings of algorithm parameters) could influence the performance of the algorithms, we found essentially no difference in the quality of solutions found by implementations of the same algorithms by different packages. However, the algorithms and even implementations of the same algorithm differ in the extent to which their execution can be parallelized, and this can have a large impact on the runtime of the algorithms, especially on highly parallel architectures. All algorithms of Inspyred and BluePyOpt, and some algorithms of Pygmo support the parallel evaluation of multiple candidate models (typically those within a particular generation or iteration), and Neuroptimus allows users to take advantage of these capabilities. It is worth noting in this context that Pygmo contains two variants of particle swarm optimization: PSO, which is closer to the original formulation of this algorithm, updates the velocities and positions of particles in a serial manner, and is thus not suitable for parallelization; and the generational variant PSOG, where the velocities and positions of all particles are updated in parallel. PSOG shares this feature with the implementation of PSO by the Inspyred package, and both of these can be run in a parallelized fashion from Neuroptimus. As a result, although all three variants of PSO produced similar final results in our tests, the runtimes of Pygmo’s PSOG and Inspyred’s PSO were significantly lower than those of Pygmo’s PSO when multiple CPU cores were utilized.

The situation is similar for the CMAES algorithm, where the current implementation in Pygmo does not support parallel evaluations. This was the reason for including the Cmaes package in Neuroptimus: this module implements CMAES in a way that allows straightforward parallelization, and Neuroptimus uses this implementation to support parallel execution of this popular and efficient method. Finally, to demonstrate the importance of parallelized implementations, we note that running a single optimization of our most complex use case (the detailed CA1 pyramidal neuron) with 10,000 model evaluations took approximately 10 days on our compute server for algorithms without parallelization; by contrast, a single run of this use case using the same number of model evaluations (e.g., 100 generations with populations of 100 individuals) with algorithms that support parallel evaluations took only a few hours on a single node of a supercomputer (accessed via the Neuroscience Gateway) that allowed an entire generation of models to be evaluated in parallel.

Our findings regarding the relative performance of various optimization methods are mostly in line with results of earlier studies that included such comparisons. Vanier and Bower [17] compared four different algorithms on a set of use cases similar to ours. They examined the performance of random search, conjugate gradient descent (a local search method), an evolutionary (or genetic) algorithm, and simulated annealing. They found that their evolutionary algorithm (which was similar to the CEO algorithm in our benchmark) delivered good performance even for more complex use cases with a larger number of parameters. This is consistent with the generally good results of evolutionary-type algorithms in our study, although we found several more recent variants that outperformed the classic version. They also found simulated annealing to be very effective, and this was later confirmed by Friedrich et al. [15] using the implementation that is built into the GENESIS simulator. None of the packages currently supported by Neuroptimus contain the traditional simulated annealing algorithm. In fact, older versions of the SciPy module used to include simulated annealing, but it was later deprecated and replaced by the basinhopping algorithm, which is considered to be a generalization of simulated annealing. In this light, the generally poor performance of the basinhopping algorithm in our tests is slightly surprising, although it may be caused by implementational problems or improper default settings of the parameters.

Our finding that CMAES performs well in a variety of different tasks is supported by several other studies. In particular, CMAES and IBEA have been compared on data-driven neuronal models, and CMAES generally delivered better final scores [41]. CMAES was also found to be efficient and robust in a study that involved fitting the biophysical parameters of models of striatal neurons [42]. Outside the neuronal modeling domain, a recent study compared two sophisticated evolution strategy variants, CMAES and xNES on different problems, and the results clearly showed that CMAES consistently outperformed xNES [43]. Our findings also support this conclusion, and add some evidence regarding two additional evolution strategy types: classic evolution strategy (CES, from the Inspyred package), which performed similarly to xNES (from Pygmo), and Pareto-archived evolution strategy (PAES, from Inspyred), which was one of the weakest performers (worse than random search) in our comparison.

### Limitations of the benchmarking study

The specific results that we obtained in our benchmarking study depend, to some extent, on some arbitrary choices that we had to take when designing our tests. For instance, we arbitrarily set the number of model evaluations to 10,000 for every algorithm to ensure a fair comparison of the final results. However, not all algorithms converged completely after 10,000 model evaluations in some of our use cases, and thus allowing more (or fewer) evaluations would likely affect the rankings based on the final score. The other performance measure that we used, the convergence score, is expected to be less sensitive to the exact number of model evaluations allowed, and also provides an indication of the speed of convergence.

We made another arbitrary choice for every algorithm with a hierarchical design, which includes all population-based methods (such as evolutionary and swarm intelligence algorithms) but also other nested algorithms such as basinhopping. We set the number of model evaluations in the innermost loop (i.e., the size of the population, or the number of steps in the local optimization) to 100, and the number of iterations in the outer loop (e.g., the number of generations) also to 100. This 100x100 partitioning of the total of 10,000 evaluations is a reasonable choice for most algorithms, and led to good results in most cases; however, it is entirely possible that a different choice (such as 200 generations with a population size of 50, or the other way around) would have resulted in improved performance for some of the algorithms.

Similarly, almost all the algorithms that we used include some tunable parameters that change the course of the optimization and may heavily influence the quality of the results. We decided to use the default settings specified in the optimization packages for all algorithms (with the exception of the CEO algorithm from the Inspyred package - see the description of this algorithm in Methods for details). In many cases, these settings were compatible with parameters recommended in the relevant literature; in some other cases, different sources suggested different settings; and in some cases, no such recommendations could be found. It is also known that the best settings for such algorithmic parameters can depend on the details of the problem, so it is possible that distinct settings would be optimal for the various use cases.

Overall, we may conclude that the rankings that we provide are not just about the algorithms themselves (or even about their implementations by particular packages), but are likely also influenced by the settings of the parameters (including the population size) for each method. In fact, we expect that the results of some algorithms could be improved substantially by using different settings, and there are also many additional optimization algorithms that were not included in the current comparison. Therefore, we hope to update and extend our study as more optimization results become available (see below for further discussion).

### Possible extensions

In addition to testing our use cases with more algorithms and settings as described above, the results of our study could be extended in other important ways. One possible direction would be to extend the set of use cases to other types of problems. All of the use cases included in the current study involved single cell models; however, in principle, Neuroptimus can also handle neural models at the subcellular and network levels, and supports the optimization of their parameters. For example, in one project, we used Neuroptimus to tune the concentrations of molecules in biochemical pathways involved in synaptic plasticity, relying on the reaction-diffusion (rxd) module of NEURON to run the simulations. In another project, we used Neuroptimus to tune synaptic weights in a network model that was constructed and simulated using Brian as an external simulator. The main reason why we did not include these examples in the current comparison is that, in both of these cases, every single simulation of the model requires more substantial resources and time, so that repeatedly running these problems with all the algorithms would have been beyond our current computational resources, and would have been completely unfeasible with algorithms that do not support parallel model evaluations.

There are also many useful features that we could potentially add to our optimization tool, Neuroptimus. For example, Neuroptimus currently returns only a single parameter combination corresponding to the lowest error score at the end of the optimization (although the parameters and errors of all the models tested are also saved into a file and may be analyzed outside Neuroptimus). This is the case even when multi-objective methods are used; the winner in this case is selected from the final population by minimizing the weighted sum of the objectives, using weights provided by the user before the optimization run. However, the final population of multi-objective optimization carries much more information, as it approximates the Pareto front (the parameter combinations representing the best possible tradeoffs between the objectives for different choices of the weights). Therefore, it would be useful to add to Neuroptimus the capability of properly representing and analyzing the results of multi-objective optimization. Other useful extensions could include the ability to chain optimization algorithms (e.g., by automatically running local optimization using the output of a global algorithm as the starting point), and the ability to visualize the progress of optimization (in error space and in the parameter space) while it is still running.

### Community and cooperation through the Neuroptimus website

We do not see the benchmarking results presented in this paper as the final word in evaluating parameter search methods for neuroscientific problems. As we argued above, it will be important to extend our study with more use cases and further evaluation of different algorithms and settings. Global parameter optimization is also a fast-moving field where new methods emerge regularly; the fact that Neuroptimus supports several actively developed packages, and is also flexible enough to accommodate new packages guarantees that new developments can be integrated with minimal effort.

We have developed and deployed the Neuroptimus web server to provide a platform for sharing and analyzing optimization results. By allowing all users to upload results obtained by using Neuroptimus, and to compare them with already uploaded results (including all the results of the current paper), the web site will become a continuously updated “live” version of this paper. This should facilitate meaningful, quantitative comparisons of parameter optimization methods, and aid the collaboration of different research groups that are interested in this topic. We encourage all interested professionals (and especially those who are experts in using particular algorithms) to run the use cases with improved settings, try other algorithms, add new use cases, and share their results on the Neuroptimus website. This way, we can collectively track new developments, and offer reliable solutions for an increasing variety of neural optimization problems.

## Methods

### Software tools and services

#### Neural Optimization User Interface (Neuroptimus)

At the core of our methodology is a software tool that we developed, called Neural Optimization User Interface (or Neuroptimus). Neuroptimus implements a software framework that allows users to set up and solve parameter optimization problems and analyze the results. Neuroptimus performs parameter optimization mainly by providing a common interface to a large number of popular parameter search algorithms implemented by various open source packages. In principle, Neuroptimus can be used to optimize the parameters of all kinds of systems; however, its main purpose is to aid parameter fitting in neural systems, and especially in detailed models of neurons. Accordingly, it includes many features that were developed specifically for this scenario, which support simulating biophysical models of neurons using the NEURON simulator, and comparing their behavior to experimental data obtained with common electrophysiological protocols.

Neuroptimus is essentially an updated and extended version of our previous tool Optimizer (https://github.com/KaliLab/optimizer) [15]. The basic design of these two pieces of software is quite similar, and they also share many details of their implementation. Therefore, we will focus on the new features and other differences here, and summarize the features that are used by the current benchmarking study, but we refer the reader to Friedrich et al. (2014) and the Neuroptimus documentation (https://neuroptimus.readthedocs.io/) for further details.

Neuroptimus is open source software, implemented in Python3, and can be accessed at the GitHub repository https://github.com/KaliLab/neuroptimus. Its functions are available both via a graphical user interface (GUI) that guides users through the steps of setting up, running, and evaluating the results of parameter optimization tasks, and via a command line interface that performs these tasks based on the settings stored in a configuration file. The GUI was built using the PyQt5 package that provides a Python binding to the cross-platform GUI toolkit Qt (version 5).

The complete definition of a neural parameter optimization problem requires the specification of multiple components. First, we need to provide the model whose parameters we wish to optimize. Neuroptimus can load, manipulate and execute models implemented in the HOC language of the NEURON simulator. The parameters to be optimized can be selected from the parameters of this model, or the user can provide a function (implemented in Python) that defines abstract parameters and how these should be mapped onto the concrete parameters of the NEURON model. As an alternative, models can be implemented by any external program that is capable of reading the variable parameters of a model candidate from a text file, setting up the model accordingly, running the simulation(s), and saving the results to files that can be interpreted by Neuroptimus.

Second, the cost function for neural parameter optimization is typically defined in terms of some target data (from experiments or prior simulations) and a function (or set of functions) that quantifies the difference between the output of the model and the target data. Neuroptimus can handle different types of target data, including time series (such as voltage and current traces), explicit spike times, and feature statistics.

Neuroptimus implements several error functions that can be used individually or in combination to evaluate during the optimization process the discrepancy between the voltage traces (or other time series) generated by the optimized model and the target data [15]. These cost functions range from general ones such as the mean squared error to more specific ones that are useful mainly in the context of fitting neuronal voltage responses and characterize the pattern and shape of action potentials (Table 1).

**Table 1:** Cost functions implemented in Neuroptimus.

| Feature name | Definition |
| --- | --- |
| Mean squared error | Mean squared difference between the model trace and the target trace point by point, normalized by the squared range of the target data |
| Mean squared error (excluding spikes) | Same as above but excludes the parts of both traces in the vicinity of action potentials (in either trace) |
| Derivative difference | Normalized mean squared difference of the temporal derivatives of the two traces |
| Spike count | Absolute difference of the number of spikes in the entire traces, normalized by the sum of the two spike counts |
| Spike count<br>(during stimulus) | Identical to spike count, except it only takes into account the action potentials during the stimulus |
| ISI difference | Sum of the absolute differences of the inter-spike intervals of the two traces, normalized by the length of the traces |
| Latency to 1st spike | Squared difference between the time to the first spike from the start of the stimulus in the two traces, normalized by the squared length of the traces |
| AP overshoot | First calculates the amplitudes of the action potentials in both traces as the difference between the AP peak voltage and the AP threshold, then takes the mean squared difference of the AP amplitudes normalized by the squared maximal amplitude of the target trace |
| AP width | Mean squared difference between the width of the action potentials at their base (at the threshold voltage level), normalized by the squared mean width of the APs in the target trace |
| AHP depth | The squared mean of the difference in the corresponding after-hyperpolarization depths, normalized by the squared range of subthreshold potential in the target trace |

The error functions above (which were already present in Optimizer; [15]) compare each voltage (or current) trace generated by a model with a specific voltage (or current) trace in the target data. However, a common task in single cell modeling involves finding model parameters such that the behavior of the model becomes similar to the typical behavior within a set of experimentally recorded neurons [5, 7]. In this case, it is more natural to define the target of the optimization as the mean values of a set of pre-selected features extracted from the experimental voltage traces (which may come from several experiments involving the same or different neurons). Then the natural way of defining error functions is by evaluating the difference between the value of a particular feature extracted from the voltage response of the model and the mean value of the same feature in the experiments, divided by the standard deviation of the feature in the experimental data. One additional advantage of this definition is that it provides standardized, dimensionless error scores that may be combined in a straightforward manner.

This approach based on feature statistics is now supported by Neuroptimus. To provide access to a diverse array of electrophysiological features, and ensure compatibility with some common workflows [5, 7, 44, 45], Neuroptimus utilizes the Electrophys Feature Extraction Library (eFEL; https://github.com/BlueBrain/eFEL) [27] to characterize the voltage responses of the models. The target data in this case contain the experimental mean and standard deviation values of a predefined set of eFEL features extracted from voltage responses to specific current step inputs, stored in a JSON file created from the recordings using the BluePyEfe tool (https://github.com/BlueBrain/BluePyEfe) and a custom script that converts the output of BluePyEfe to the format expected by Neuroptimus. This JSON file also contains the full specification of the stimulation protocols. When the optimization is run using the GUI, the settings of the stimuli and the features are automatically loaded into the GUI from this input file. During the optimization process, in every model evaluation step, the features included in the input file (and selected in the GUI) are extracted from the model’s voltage traces, and errors are computed for every feature using the feature statistics-based error function described above [46].

Recent studies classify optimization problems according to the cardinality of objectives as single, multi- (2-3 objectives) and many-objective tasks (more than 3 dimensions), which affects the nature and the complexity of the problem [47]. However, we characterized our problems simply as single- or multi-objective problems because these require different internal representations and are solved by different algorithms. Multi-objective problems involve several objective functions that are to be minimized simultaneously and require finding a set of solutions that give the best tradeoffs between the objectives.

Neuroptimus makes it possible to use arbitrary weighted sums of error functions as the ultimate objective function of the parameter search. When single-objective algorithms are used, the weighted sum is calculated for every model during the optimization process, and is used as the objective function. In the case of multi-objective algorithms, all the error functions are treated as separate objectives during the optimization, but the weighted sum is still used after running the search to select a single preferred solution from those returned by the algorithm [5, 23].

Neuroptimus supports parameter optimization algorithms implemented by five external Python packages (Pygmo, Inspyred, BluePyOpt, Scipy, and Cmaes), and also contains an internal implementation of a simple random search algorithm that takes independent, uniformly distributed samples from the entire search space. Pygmo is a general-purpose scientific Python library for optimization, based on the C++ library pagmo, which implements many different optimization algorithms in a common framework [48]. Inspyred is a Python library specifically developed for bio-inspired (mainly evolutionary) computation, and was already supported by Optimizer [15]. The Blue Brain Python Optimization Library (BluePyOpt) is a software framework developed at the Swiss Blue Brain Project [23], which implements multi-objective optimization algorithms including the Indicator Based Evolutionary Algorithm (IBEA), and has been applied successfully in several computational neuroscience projects [5, 7, 44, 49, 50, 51, 52, 53]. SciPy [54] provides implementations of various methods for scientific computation, and includes several basic optimization algorithms, some of which were already supported by Optimizer [15]. Finally, we also included the Cmaes package because it provides a simple, robust, and easily parallelizable implementation of the Covariance Matrix Adaptation Evolution Strategy (CMAES) algorithm, a popular and powerful search method that is also included in Pygmo but in an implementation that does not support the parallel evaluation of models within a population.

Some of the algorithms are local (essentially gradient-based) search methods, but most of them are based on metaheuristics that attempt to find the global minimum of the cost function(s). Many of the most popular single- and multi-objective optimization algorithms are included. Most of the algorithms also have parameters that are configurable through the GUI or the configuration file.

Solving nontrivial parameter optimization problems typically requires the evaluation of many parameter combinations. In our case, this corresponds to running a large number of simulations, which may take a prohibitively long time if simulations are performed sequentially, especially for complex models such as morphologically detailed neurons, circuits, or multi- scale models that include biochemical or molecular processes. Fortunately, many global optimization methods (including evolutionary and swarm intelligence algorithms) can be implemented in a way that populations of models can be evaluated in parallel, and several such parallel (or easily parallelizable) implementations are included in the Python libraries supported by Neuroptimus. However, Python provides several different methods for parallel execution of code, and the optimization packages we use differ in terms of which parallelization approaches they support. As a consequence, Neuroptimus uses the *multiprocessing* module for the parallel execution of algorithms in Pygmo, Inspyred and Cmaes, while it relies on the IPython Parallel (*ipyparallel*) package to run the algorithms of BluePyOpt in parallel. We note that some optimization algorithms cannot be efficiently parallelized, while for some others (including several in the Pygmo package) parallel execution is not currently supported by the optimization library.

Batch evaluation of the models is a requisite to use one of the various parallelization strategies in the Neuroptimus. Therefore both internal and external evaluations have to generate results simultaneously. If we used a single model instance in every process, the results could be mixed or swapped. Therefore, when simulations are carried out within Neuroptimus (using NEURON), a new model instance is created for every parameter set generated by the selected algorithm, and every evaluation running in parallel is performed with a separate model. In case of using the external simulator, every individual is evaluated in a separate subprocess, and files with unique names are used for communication between Neuroptimus and the external simulation script.

We list all of the available algorithms along with their basic properties in Table 2. Many of these algorithms were tested in our benchmarking study, and these will be described in more detail below.

**Table 2:**
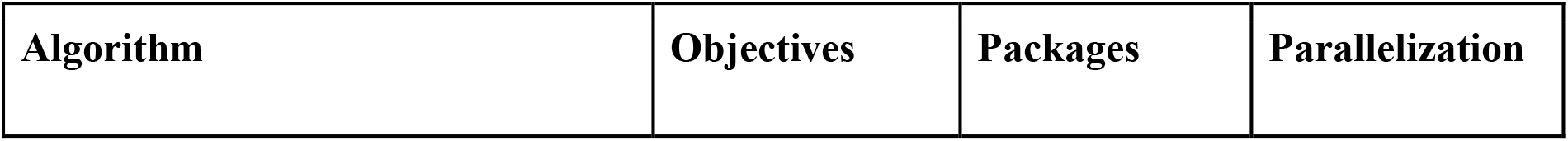
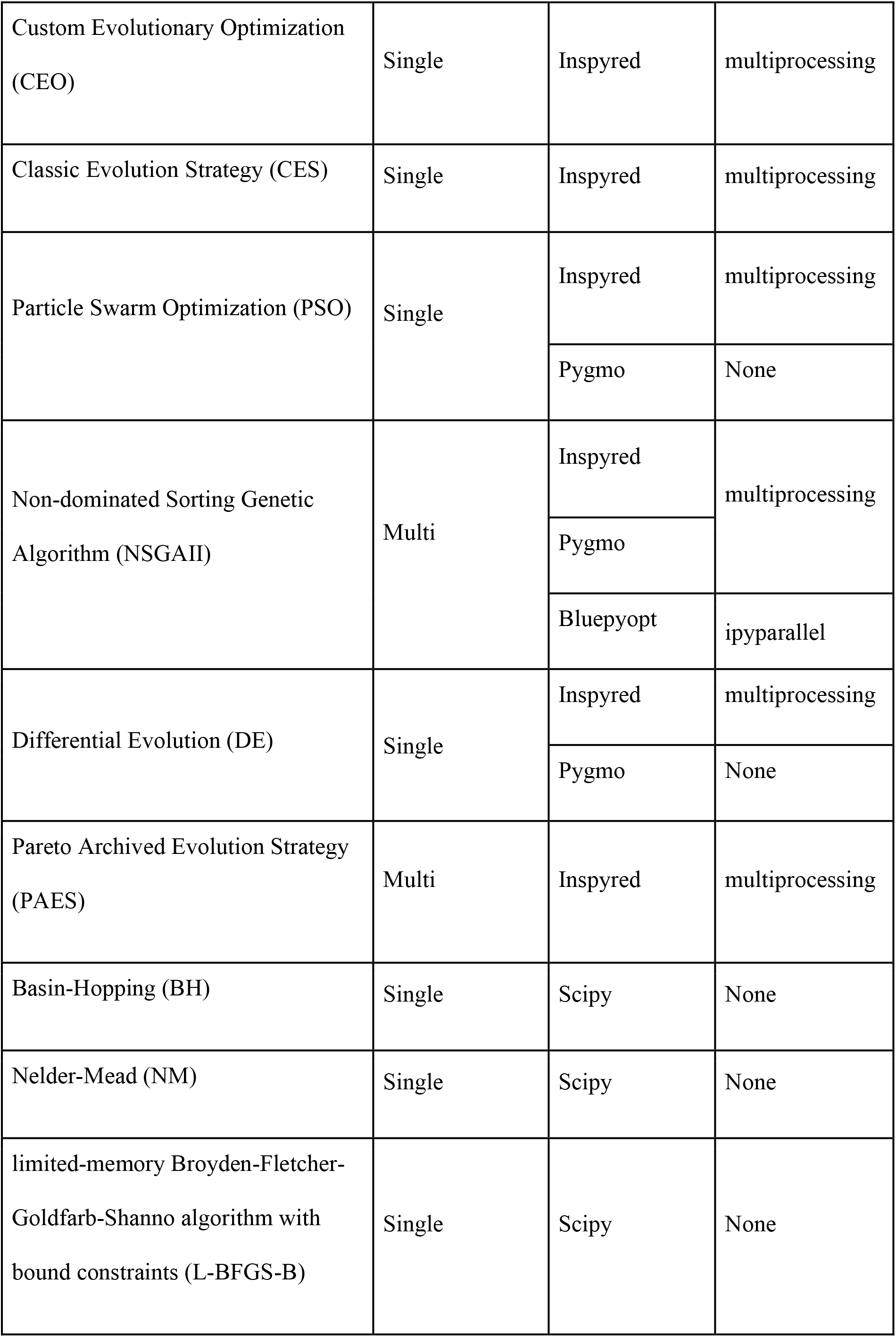
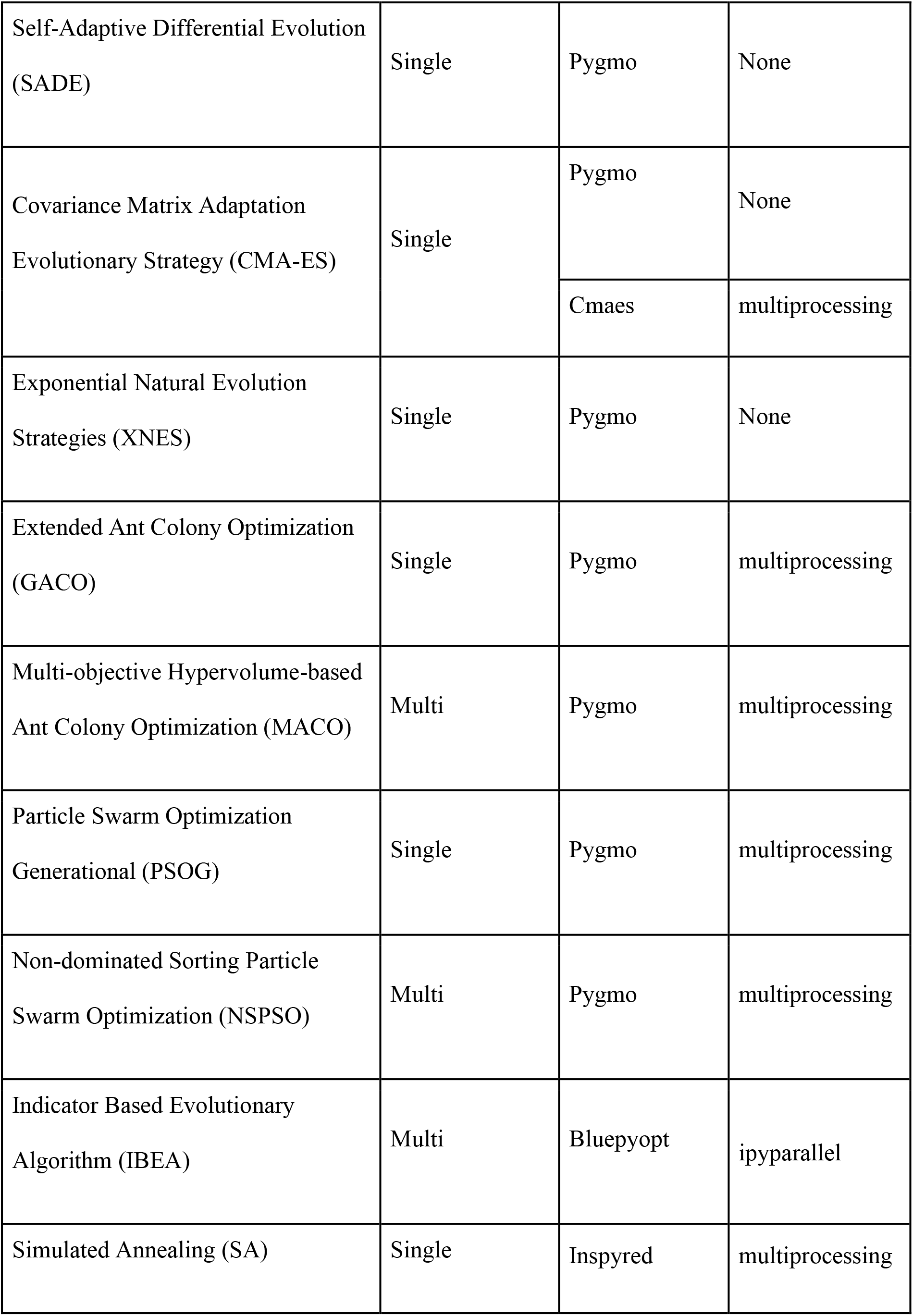
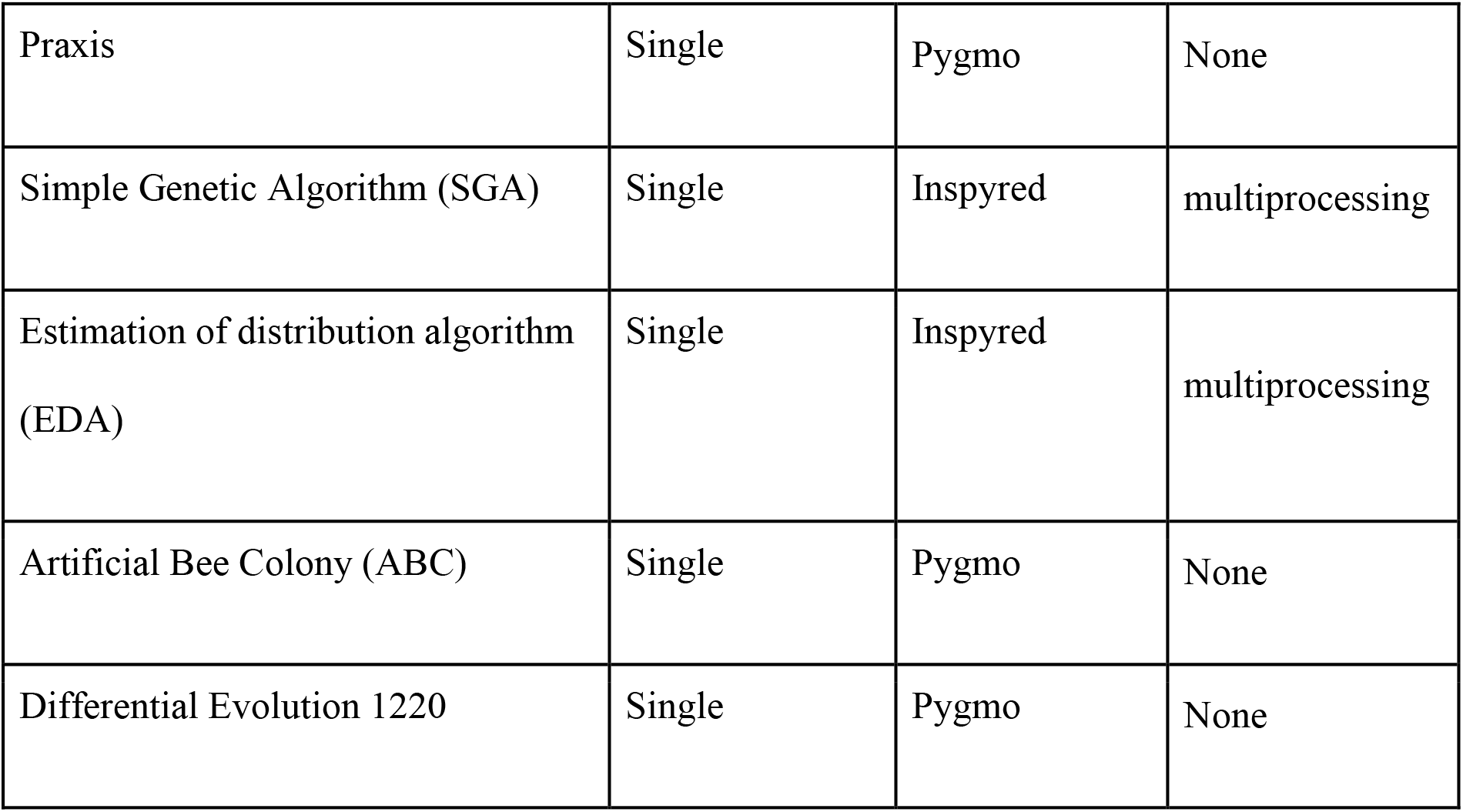
Algorithms included in Neuroptimus. The properties listed include the full name of the algorithm, the abbreviation used in this article, the type according to the number of objectives (single/multi-objective), the implementing package(s), and the method of parallelization used in Neuroptimus (None if only serial execution is supported).

The easiest way to perform parameter optimization using Neuroptimus is by using the GUI, whose seven tabs guide the user through the steps of setting up, running, and evaluating the results of the parameter search. The GUI allows the user to load the target data, select the model and the parameters to be optimized, set up the stimulation and recording conditions, configure the error function(s), run the parameter search, and then visualize and analyze the results. The final as well as intermediate results of the optimization are also saved to files, and can be analyzed outside Neuroptimus. This includes the parameters and errors of each simulated model as well as the statistics of generations saved into text files, the voltage trace of the best model saved into text files and in several image formats, and a final summary of the optimization process and the results saved into an HTML file for visual inspection through a web browser and a JSON file for a machine-readable non-SQL data representation. This final metadata file created after the optimization contains automatically generated names for the optimization and the model, details of the parameters of the model (name, boundaries and optimal values), details of the error functions used to calculate the final error (name, value, weight, weighted value), settings of the target data, the algorithm and package used for the optimization, parameters used by the algorithm, and finally the statistics of each generation.

The program also saves the full configuration of the optimization task, and the resulting configuration file can be used (directly, or after suitable modifications) by the command-line interface of Neuroptimus to re-run the optimization (with the same or modified settings). This method was used in our benchmarking study to run batches of the same optimization with different random seeds, using a simple Python script to edit the configuration file and create multiple versions of the optimization task.

#### Neuroptimus server

To share our results in a way that allows easy replication and further analysis, and to enable the straightforward extension and updating of the current study, we created an online database of optimization results that is accessible via a web interface. We designed, created and deployed the Neuroptimus web-server, which can be publicly accessed at https://neuroptimus.koki.hu and enables all users to browse, view and analyze the optimization results stored in the database. Furthermore, authenticated users can also upload their optimizations and compare their results with previously uploaded ones.

The Neuroptimus server structure consists of an Nginx web server that handles the requests and responses, the frontend implemented using the JavaScript library ReactJS, the backend created in the Python web framework Django, backed up by a PostgreSQL database connection. The site handles the authentication of registered users, uploading of optimization results via a web form, visualization of the data in a table structure, and the creation of plots for comparison. The database stores information about the optimization itself, the model used for the optimization and its parameters, the algorithm and its configuration, details about the target data, the statistics of each generation produced by the algorithm, creation time of results and upload time. The metadata JSON file created by Neuroptimus can be uploaded to the server and all of its information content is transferred to the database automatically. Optionally the compressed optimization files can also be uploaded and subsequently downloaded. Analysis of the optimization can be created semi-automatically by selecting the desired algorithms for comparison and visualizing them on the charts. Thus far generation plots, final and convergence score box plots are available for online observation.

During the registration process users need to provide their name, affiliation, and email address, choose a username, and create a password. Verifying email addresses grants permission for users to upload their optimizations. Forgotten passwords can be reset on the website via email verification.

### Optimization algorithms

In the current study, we evaluated a large set of parameter search algorithms, including several of the most widely used single-objective and multi-objective methods. Our optimization tool supports optimization algorithms implemented by five separate Python packages: Inspyred [55], Pygmo [48], BluePyOpt [23], Cmaes [56], and Scipy [54]. Table 2 shows which packages implement each of the supported algorithms. The majority of these algorithms can be categorized as evolutionary or nature-inspired metaheuristics.

Due to constraints on time and computational resources, we could not include every single algorithm supported by Neuroptimus in the detailed comparison that we performed using our neural benchmarking suite (see below). However, we aimed to provide good coverage of algorithms that were used previously in neuronal optimization [9, 15, 17, 23], and also included several additional algorithms that consistently provided good performance in other settings [57, 58, 59, 60].

Finally, we added some basic search methods such as uniform random sampling and two widely used local optimization algorithms to provide a baseline against which we can measure the performance of more sophisticated methods. The following algorithms were tested in our neural optimization benchmark.

#### Baseline algorithm

The *Random Search (RAND)* algorithm is the simplest heuristic to discover solutions by trial and error. This is our baseline method, which samples parameters from the search space repeatedly based on the uniform probability distribution. Neuroptimus uses our own implementation of this method [61].

#### Local optimization algorithms

The *Nelder-Mead (NM)* algorithm is a classic simplex-based direct search method to find a local minimum of the cost function [62].

The *limited-memory Broyden-Fletcher-Goldfarb-Shanno algorithm with bound constraints (L-BFGS-B)* is considered to be a modern and efficient algorithm that aims to find a local minimum of the objective function using a limited amount of computer memory [63].

#### Single-objective global optimization algorithms

The *Custom Evolutionary Optimization (CEO)* algorithm is a relatively simple member of the large class of evolutionary optimization algorithms. Evolutionary algorithms are metaheuristics for global optimization inspired by biological evolution. Each candidate solution, represented by a particular combination of the unknown parameters, is considered to be an individual within a population, and the value of the cost function for that parameter combination is treated as the “fitness” of that individual (with lower costs normally associated with higher fitness). The initial population typically consists of random samples from the search space. The population is then updated through the application of various operators. New individuals are generated via the application of genetic operators such as mutation, which introduces random variations into the parameters of an individual, and crossover, which randomly combines the parameters of two individuals. The size of the population is maintained by selecting individuals with higher fitness. These steps are repeated iteratively for a certain number of generations. Many different variants of evolutionary algorithms exist that differ in the details of the operators, and may also apply additional heuristics. The CEO algorithm is based on the EvolutionaryComputation class of the Inspyred package, and uses Gaussian mutation and blend crossover variators.

The *Classic Evolution Strategy (CES)* algorithm belongs to a subclass of evolutionary optimization algorithms called evolution strategies. In these algorithms, there are distinct mutation rates associated with each parameter, and these mutation rates are changed adaptively during the optimization [64].

The *Covariance Matrix Adaptation Evolution Strategy (CMA-ES)* algorithm is an evolutionary algorithm which samples candidate solutions from multivariate normal distributions with adapting mean and covariance matrix [65].

The *Exponential Natural Evolution Strategy (XNES)* algorithm is an evolution strategy (ES) that uses the natural gradient to update the search distribution [66].

The *Differential Evolution (DE)* algorithm is an evolutionary algorithm that generates new candidate solutions from existing individuals based on some simple mathematical rules [57].

The *Self-Adaptive Differential Evolution (SADE)* algorithm is a version of the Differential Evolution algorithm, which adjusts the mutation rate and the crossover rate adaptively [67].

The *Particle Swarm Optimization (PSO)* algorithm represents candidate solutions as particles moving around in the search space. Each particle has a velocity and moves by adding this to its current position in every iteration. Initially the velocity is random, and it is modified after each iteration, influenced by the currently known best positions for the individual particles and that of the entire group. In this basic implementation, velocity and position updates are carried out sequentially for each particle [68].

The *Particle Swarm Optimization Generational (PSOG)* algorithm is similar to the PSO algorithm above but, in every iteration, it first updates the velocity for all particles, then updates the positions. This allows efficient parallel execution of the algorithm.

The *Extended Ant Colony Optimization (GACO)* algorithm is a bio-inspired algorithm based on the analogy of ants finding paths from colony to food. In this algorithm, artificial agents move through the parameter space, and lay down “pheromones” depending on the quality of the solutions they find. These pheromones attract the other agents, making it more likely that they move to locations with high amounts of pheromone. This extended version of the algorithm calculates the locations of future generations of ants by sampling from a multi-kernel Gaussian distribution that depends on the quality of previously found solutions [69].

The *Basin-Hopping (BH)* algorithm is a generalization of the Simulated Annealing algorithm that was used in several earlier studies of neural parameter optimization [15, 17]. Basin-hopping is a two-level algorithm: its outer loop performs stochastic jumps in the search space, while the inner loop performs local optimization. The resulting new local minimum is always accepted if it is better than the previous one, but it may also be accepted if it is worse with a probability that depends on the increase in the cost function [70].

#### Multi-objective global optimization algorithms

The *Non-dominated Sorting Genetic Algorithm II (NSGA2)* is an evolutionary multi-objective algorithm. Multi-objective optimization algorithms aim to optimize several cost functions simultaneously, trying to find non-dominated (or Pareto-optimal) solutions where none of the cost functions can be improved without degrading the performance on some other cost functions. The algorithms also aim to create a diverse set of solutions that collectively provide good coverage of the Pareto front. In NSGA2, a child population is created from the parent population using the usual genetic operators, mutation and crossover. Individuals in the next generation are then selected from the joint population based on Pareto dominance and the so-called crowding distance that penalizes closely related individuals and helps maintain diversity within the population [58].

The *Pareto Archived Evolution Strategy (PAES)* algorithm is a simple multi-objective algorithm that uses local search (mutation) from the current individual(s) and maintains a reference archive of previously found non-dominated solutions to approximate the dominance ranking of candidate solutions [71].

The *Indicator Based Evolutionary Algorithm (IBEA)* is a multi-objective evolutionary algorithm that computes the fitness value based on predefined binary indicators. It performs environmental selection by removing the worst individuals, chooses parents by comparing the fitness values of randomly selected pairs of individuals, and applies mutation and crossover to create offspring, repeating the process iteratively until reaching the maximum number of generations [72].

The *Multi-objective Hypervolume-based Ant Colony Optimizer (MACO)* is a multi-objective optimization algorithm that extends the GACO algorithm described above, combining hypervolume computation and non-dominated fronts for ranking individuals [73].

The *Non-dominated Sorting PSO (NSPSO)* algorithm extends PSO by making a better use of personal bests and offspring for non-dominated comparison [74].

### Use cases

To compare the efficiency of various parameter search methods in solving neuronal parameter optimization tasks, we designed and implemented a suite of six different problems that may be considered typical use cases in this domain. All of these use cases can be handled by Neuroptimus, which allowed us to run all benchmarks using every selected algorithm within the same framework, and made the subsequent evaluation of their performance quite straightforward (see below). Five of the use cases were similar (or identical) to those presented in [15], although some of them were modified to increase the robustness of the simulations (avoiding errors due to invalid parameter combinations, in the case of the AdExpIF example) or to move the target behavior of the model away from a critical boundary (the transition to repetitive firing, in the case of the Hodgkin-Huxley model). We provide a description of each use case below; all the files required to run these examples, along with detailed guides to setting up the optimizations in the Neuroptimus GUI, can be found in the corresponding subfolders of the neuroptimus/new_test_files directory of the Neuroptimus Github repository (https://github.com/KaliLab/neuroptimus).

#### Hodgkin-Huxley

This use case is based on a single-compartment model, which contains conductances from the original Hodgkin-Huxley model (Na+, K+, leak )[28], and is implemented in NEURON. To generate the target voltage trace, a suprathreshold step current was injected into the soma of the neuron model (amplitude = 300 pA, delay = 200 ms, duration = 500 ms, and the voltage trace duration is 1000 ms). The test case involves recovering the correct conductance densities (3 parameters) that were used to generate the target trace, while keeping the properties of the currents and the other parameters of the model constant (at their original value). A combination of four features (spike count, spike amplitude, spike width, mean squared error of voltage excluding spikes) was used to compare each simulated trace to the original (target) trace.

#### Voltage Clamp

In the Voltage Clamp benchmark problem the same single-compartment model with the same conductances is used as in the Hodgkin-Huxley problem. In addition, this model contains a conductance-based synapse. The goal here is to recover the synaptic parameters (weight, rise and decay time constants, delay – 4 parameters) from simulated voltage clamp recordings during synaptic stimulation (four presynaptic spikes at 10 Hz), using the mean squared error cost function to compare the current traces.

#### Passive, anatomically detailed neuron

This benchmark uses a morphologically detailed passive model of a hippocampal CA1 pyramidal cell implemented in NEURON. During the experiment, a short (3 ms, 500 pA) and a long (600 ms, 10 pA) current pulse (separated by 300 ms) were injected into the soma, and the membrane potential was also recorded there. The neuron was filled with a dye during the recording, and was reconstructed using Neurolucida. This reconstruction defines the morphology of the model, and the task involves fitting 3 passive parameters (specific capacitance, leak conductance density, specific axial resistance, all of which are assumed to be uniform within the cell) to reproduce the experimental data recorded using the same complex current clamp stimulus. Traces are compared via the mean squared error cost function. All the experimental data for this use case, including the morphological reconstruction and the electrophysiological recordings, were provided by Miklós Szoboszlay and Zoltán Nusser.

#### Simplified model

This use case attempts to fit the behavior of a six-compartmental simplification of a biophysically accurate and morphologically detailed hippocampal CA1 pyramidal cell model [35] to the somatic voltage responses of the original model with full morphology. Both models contained the same set of voltage-gated conductances in their somatic and dendritic compartments: transient Na channels (separate somatic and dendritic subtypes), delayed rectifier, A-type, and M-type voltage-gated K channels, C-type and AHP-associated Ca- dependent K channels, L-type and N-type Ca channels, and the hyperpolarization-activated HCN channels. Dendrites of the full model were clustered based on their passive voltage responses, and each of these clusters defined a dendritic compartment in the simplified model. The densities of ion channels in the dendritic compartments of the simplified model were set to the average values in the corresponding clusters of the full model, while the densities of the nine somatic conductances were subject to parameter optimization. The original full model was implemented in GENESIS, while the simplified model was implemented in the NEURON simulator. The target data was the voltage response of the full model to the injection of a 200 pA step current stimulus into the soma (the stimulus started at 200 ms and lasted for 600 ms, with a total recording duration of 1000 ms). The fit was evaluated via a combination of features including mean squared error (excluding spikes) weighted by 0.2, spike count (weight 0.4), latency to first spike (weight 0.1), action potential amplitude (weight 0.1), action potential width (weight 0.1), and after-hyperpolarization depth (weight 0.1).

#### Extended integrate-and-fire model

In this benchmark problem, the parameters of a phenomenological (adaptive exponential integrate-and-fire) spiking model [32, 33], implemented in the NEST simulator [34] were fitted to capture the somatic responses of a real neuron (hippocampal CA3 pyramidal cell) to four different inputs. Voltage traces were recorded experimentally in response to current steps of 900 ms duration, and 0.30, 0.35, 0.40, and 0.45 nA amplitudes (the step was delayed by 100 ms, and the recordings lasted for 1100 ms). Sampling frequency was 5 kHz. The unknown parameters to be optimized were the capacitance, the leak conductance, the reversal potential of the leak current, the threshold voltage, the reset voltage, the refractory period, the steepness of the exponential part of the current-voltage relation, the subthreshold adaptation conductance, the spike adaptation current, and the adaptation time constant (10 parameters). During the optimization the mean squared error (excluding spikes), the spike count (during stimulus), and the latency to first spike error functions were used with equal weights.

#### CA1 pyramidal cell

This is our most complex benchmark problem both regarding the number of parameters to be optimized and the complexity of the model. The test case is based on an anatomically and biophysically detailed rat hippocampal CA1 pyramidal cell model built for the NEURON simulator in our research group. The morphology of the model was from [39]. The model contained several different voltage-gated ion channels in its somatic, dendritic, and axonal compartments: a transient Na conductance, delayed rectifier, A-type, M-type, and D-type voltage-gated K conductances, and the hyperpolarization-activated current Ih. Many attributes of the model were well-constrained by experimental observations available in the literature, including the distributions and kinetic properties of the ion channels. The target data (provided by Judit Makara) consisted of the means (and associated standard deviations) of 20 different types of features extracted by eFEL from the voltage responses of five CA1 pyramidal neurons to somatic current step injections of six different amplitudes (-0.25, 0.05, 0.1, 0.15, 0.2 0.25 nA), with each stimulus repeated three times for every cell. This resulted in a total of 66 feature values to be matched by the model. The eFEL features and the associated current step amplitudes are listed in Table 3.

**Table 3:** List of eFEL features (with brief explanations) and associated current amplitudes used as the target data in the detailed CA1 pyramidal neuron use case.

| Feature name from eFEL | Feature description | Fitted current steps (nA) |
| --- | --- | --- |
| Spikecount | Number of spikes during stimulus | 0.05, 0.1, 0.15, 0.2, 0.25 |
| inv_first_ISI | Reciprocal of first interspike interval | 0.05, 0.1, 0.15, 0.2, 0.25 |
| inv_last_ISI | Reciprocal of last interspike interval | 0.05, 0.1, 0.15, 0.2, 0.25 |
| inv_time_to_first_spike | Reciprocal of time to first spike | 0.05, 0.1, 0.15, 0.2, 0.25 |
| steady_state_voltage | Average voltage after stimulus | -0.25, 0.05, 0.1, 0.15, 0.2, 0.25 |
| voltage_base | Average voltage during the last 10% of time before the stimulus | -0.25, 0.05, 0.1, 0.15, 0.2, 0.25 |
| voltage_deflection | Difference between the voltage base and the steady-state voltage at the end of the stimulus | -0.25 |
| voltage_deflection_begin | Difference between the voltage base and the mean voltage in the early phase (5 to 15% of duration) of the stimulus | -0.25 |
| AHP_depth_abs | Absolute value of the depth of afterhyperpolarization | 0.15, 0.2, 0.25 |
| AHP_time_from_peak | Time from the peak of the AP to the minimum of the afterhyperpolarization | 0.15, 0.2, 0.25 |
| AP2_amp | Amplitude of second AP | 0.15, 0.2, 0.25 |
| AP_amplitude | Average height of APs | 0.15, 0.2, 0.25 |
| AP_begin_voltage | Threshold of AP initiation | 0.15, 0.2, 0.25 |
| AP_duration_half_width | Half-width of the AP | 0.15, 0.2, 0.25 |
| AP_fall_time | Time from the peak of the AP to the end of the AP | 0.15, 0.2, 0.25 |
| AP_rise_time | Time from the AP threshold to the peak of the AP | 0.15, 0.2, 0.25 |
| AP_last_amp | Amplitude of the last AP | 0.15, 0.2, 0.25 |
| time_to_last_spike | Time to the last spike | 0.15, 0.2, 0.25 |
| sag_amplitude | Difference between the minimal voltage and the steady state voltage at the end of the stimulus | -0.25 |
| sag_ratio1 | Ratio between sag amplitude and maximal hyperpolarization from voltage base | -0.25 |

We optimized 12 abstract parameters of the model that were mapped onto the actual parameters of the NEURON implementation by an appropriate user function. Ten parameters determined the densities of the voltage-gated and leak conductances in the different compartments (soma, dendrites, axon) of the neuron; one parameter represented the reversal potential of the leak current; and the final parameter determined the difference between the half-activation and half-inactivation potential values of the Na conductance.

### Evaluation Methods

We tested the different optimization algorithms on each of the six model optimization tasks described above. To ensure a fair comparison of model performance, we allowed 10,000 model evaluations for every algorithm on each task. For all population-based methods (including evolutionary algorithms and swarm intelligence-based approaches) we set the population size to 100, and the number of generations to 100 as well. We similarly set 100 global and 100 local minimization steps for two-stage algorithms. Otherwise, we ran every algorithm with its default settings in Neuroptimus. These default options are typically the package default settings, with one significant exception: we observed that the default settings of the EvolutionaryComputation class of the Inspyred package that underlies our CEO algorithm led to essentially no optimization, so we adjusted the default number of elites from 0 to half of the population size, changed the mutation rate from 0.1 to 0.25 and standard deviation of Gaussian mutation from 1 to 0.5.

Optimization runs were parallelized for all algorithms where this is supported by Neuroptimus and the underlying packages (see Table 2). For the most resource-intensive use case (the detailed CA1 pyramidal neuron model) these parallelized runs were performed on supercomputers via the Neuroscience Gateway [75]; simpler use cases and algorithms that do not support parallelization were run on a Dell PowerEdge R730 compute server or personal computers. To allow meaningful statistical comparisons between the algorithms, we performed 10 independent runs (using distinct random seeds) of each algorithm in every use case.

We visualized and compared the performance of the algorithms in each use case using several different methods. All the comparisons were based on the change in the total error during the optimization. First, we visualized the convergence of the algorithms by plotting the cumulative minimum of the error function after every generation (i.e., after every 100 model evaluations). We plotted the median value across the 10 runs to see which algorithms typically find the best solutions after a given number of model evaluations. The lowest and the highest errors achieved by the 10 runs were also calculated in every iteration to observe how well the algorithm performs in the best case, and whether it gets stuck in some cases.

We defined two basic scores to characterize and compare the performance of the algorithms in a concise manner. The first of these scores was defined as the lowest error achieved during the entire optimization run (these are usually, but not always, associated with members of the final population). We visualized the distribution of this measure across the 10 independent runs using box plots that show the median, interquartile range, minimum, and maximum values, and also indicate apparent outliers.

In the case of more complex, detailed models, each model evaluation (simulation) can be time-consuming, and thus we are also interested in which algorithms can find a reasonably good solution in a relatively short time. To characterize the convergence speed of an algorithm, we used the sum of the logarithms of the error scores achieved by the best individuals in each generation. This is essentially the area under the logarithmic convergence curve - the smaller this sum is, the faster the algorithm found a relatively good solution.

## Notes

### Competing Interest Statement

The authors have declared no competing interest.

